# Enhanced RNAi does not provide efficient innate antiviral immunity in mice *in vivo*

**DOI:** 10.1101/2024.07.29.605661

**Authors:** Marcos Iuri Roos Kulmann, Eliska Taborska, Brigita Benköova, Martin Palus, Ales Drobek, Filip Horvat, Josef Pasulka, Radek Malik, Eva Salyova, Vaclav Hönig, Michaela Pellerova, Maria Borsanyiova, Ondrej Stepanek, Shubhada Bopegamage, Daniel Ruzek, Petr Svoboda

## Abstract

In RNA interference (RNAi), long double-stranded RNA (dsRNA) is cleaved by Dicer endonuclease into small RNA interfering RNAs (siRNAs), which guide degradation of complementary RNAs. While RNAi mediates antiviral innate immunity in plants and many invertebrates, vertebrates adopted sequence-independent response and their Dicer produces siRNAs inefficiently because it is adapted to process small hairpin microRNA precursors in the gene-regulating microRNA pathway. Mammalian RNAi is thus a rudimentary pathway of unclear significance. To investigate its antiviral potential, we modified mouse Dicer locus to express a truncated variant (Dicer^ΔHEL1^) known to stimulate RNAi. Next, we analyzed how Dicer^ΔHEL1/wt^ mice respond to four RNA viruses: Coxsackievirus B3 (CVB3) and encephalomyocarditis virus (ECMV) from *Picornaviridae*; tick-borne encephalitis virus (TBEV) from *Flaviviridae*; and lymphocytic choriomeningitis virus (LCMV) from *Arenaviridae*. Increased Dicer activity in Dicer^ΔHEL1/wt^ mice did not elicit any antiviral effect. supporting insignificant antiviral function of endogenous mammalian RNAi *in vivo*. However, we also report that sufficiently high expression of Dicer^ΔHEL1^ suppressed LCMV in embryonic stem cells and in a transgenic mouse model. Altogether, mice with increased Dicer activity offer a new benchmark for identifying and studying viruses susceptible to mammalian RNAi *in vivo*.

## Introduction

RNAi is sequence-specific RNA degradation induced by long dsRNA (Fire et al., 1998). RNAi starts with RNase III Dicer cutting long dsRNA into 21-23 nt long siRNAs, which are loaded onto Argonaute endonucleases to form RNA-induced Silencing Complex (RISC), in which they serve as guides for recognition and cleavage of complementary RNAs. RNAi acquired many biological roles including gene regulation, antiviral immunity or defense against mobile elements (reviewed in (Ketting, 2011)). At the same time, vertebrate evolution brought curtailed RNAi and adaptation of Dicer to biogenesis of gene-regulating microRNAs (miRNAs) (reviewed in (Bartel, 2018)), which are small RNAs released by Dicer from genome-encoded small hairpin precursors.

The key feature of Dicer differentiating RNAi and/or miRNA pathway support is its N-terminal helicase domain, which can mediate recognition and ATP-dependent processive cleavage of long dsRNA and/or promote ATP-independent pre-miRNA recognition and processing (reviewed in (Zapletal et al., 2023)). During vertebrate evolution, the N-terminal helicase domain lost ATPase activity and evolved to support high-fidelity processing of pre-miRNA (Aderounmu et al., 2023; Zapletal et al., 2022; Zhang et al., 2002). In fact, adaptation of mammalian Dicer for miRNA biogenesis made the N-terminal helicase an autoinhibitory element of long dsRNA processing (Ma et al., 2008). Consequently, the dominant class of Dicer products in almost all investigated mammalian cells are miRNAs. Mammals retain residual canonical RNAi response, i.e. mammalian Dicer is still able to cleave dsRNA into siRNAs, which are loaded onto AGO2, which can mediate sequence-specific endonucleolytic cleavage of perfectly complementary targets (Liu et al., 2004; Meister et al., 2004; Song et al., 2004). However, canonical RNAi in most mammalian cells is inefficient, in part owing to inefficient processing of long dsRNA into siRNAs (Demeter et al., 2019; Chakravarthy et al., 2010; Ma et al., 2008; Zhang et al., 2002) and in part to other long dsRNA responses in mammalian cells (including adenosine deamination and the interferon response), which hamper RNAi (Demeter et al., 2019; Seo et al., 2013; Takahashi et al., 2018; van der Veen et al., 2018). An exception of the rule is the mouse oocyte where functionally important endogenous RNAi emerged through an oocyte-specific promoter of retrotransposon origin, which expresses a truncated Dicer variant (denoted Dicer^O^). Dicer^O^ lacks the HEL1 subdomain of the N-terminal helicase, efficiently cleaves long dsRNA, and supports functionally relevant canonical RNAi (Demeter et al., 2019; Flemr et al., 2013).

RNAi can provide antiviral innate immunity. It was first shown for the flock house virus (FHV) (Lu et al., 2005) and the vesicular stomatitis virus (VSV) (Schott et al., 2005; Wilkins et al., 2005), viruses of broad host range, and later with natural viruses of *C. elegans* (Felix et al., 2011; Sarkies et al., 2013) and *D. melanogaster* (Saleh et al., 2009; Tassetto et al., 2017). The situation in mammals is convoluted. In 2013, two reports provided evidence suggesting that RNAi may have an antiviral role in mammals (Li et al., 2013; Maillard et al., 2013), which were immediately intensely debated (Cullen et al., 2013). Following the initial findings on Nodamura virus (NoV) and encephalomyocarditis virus (EMCV), several other studies reported functional antiviral RNAi against influenza A virus (IAV) (Li et al., 2016), human enterovirus 71 (HEV71) (Baldaccini et al., 2024; Fang et al., 2021; Qiu et al., 2017), Zika virus (ZIKV) (Xu et al., 2019; Zhang et al., 2022), dengue virus type 2 (DENV2) (Kakumani et al., 2013; Qiu et al., 2020), Semliki forest virus (SFV) (Baldaccini et al., 2024), and Sindbis virus (SINV) (Baldaccini et al., 2024; Zhang et al., 2022). Showing effects in mice and/or in cultured cells, these studies argued that i) viral siRNAs are produced in infected cells and suppress viruses, ii) antiviral effects are most visible in interferon weakened/deficient hosts, and iii) viruses evolved viral suppressor of RNAi (VSR) proteins counteracting RNAi (Fang et al., 2021; Han et al., 2020; Maillard et al., 2013; Qiu et al., 2020). At the same time, other studies showed negligible levels or absent virus-derived siRNAs (vsiRNAs) and no antiviral effects of RNAi for a number of tested viruses including IAV (Bogerd et al., 2014), DENV2 (Bogerd et al., 2014; Parameswaran et al., 2010), SINV (Bogerd et al., 2014; Girardi et al., 2013; Girardi et al., 2015; Schuster et al., 2019), hepatitis C virus (HCV) (Parameswaran et al., 2010), West Nile virus (WNV) (Bogerd et al., 2014; Parameswaran et al., 2010), yellow fever virus (YFV) (Bogerd et al., 2014; Schuster et al., 2019), poliovirus (Parameswaran et al., 2010), Venezuelan equine encephalitis virus (VEEV) (Bogerd et al., 2014), coxsackievirus B3 (CBV3) (Schuster et al., 2019; Schuster et al., 2017), vesicular stomatitis virus (VSV) (Baldaccini et al., 2024; Bogerd et al., 2014; Parameswaran et al., 2010), measles virus (Bogerd et al., 2014), herpes simplex virus type 1 (HSV-1) (Bogerd et al., 2014), SARS-CoV-2 (Baldaccini et al., 2024), and reovirus (Bogerd et al., 2014).

To obtain new insights into the antiviral potential of mammalian RNAi *in vivo*, we investigated viral resistance of mice where RNAi activity was increased by expressing a Dicer^O^-like variant (Buccheri et al., 2024; Zapletal et al., 2022). We aimed at increasing siRNA production from long dsRNA while having a minimal effect on the miRNA pathway. In the main genetic model, we modified the endogenous Dicer locus to produce an N-terminally truncated Dicer variant (Dicer^ΔHEL1^, Fig. 1A), which is functionally equivalent to naturally occurring Dicer^O^ (Zapletal et al., 2022). Although *Dicer*^Δ*HEL1/*ΔHEL1^ mice die perinatally from developmental defects, *Dicer*^Δ*HEL1/wt*^ mice are viable and fertile (Zapletal et al., 2022). Importantly, converting half of endogenous Dicer expression into the highly active Dicer variant *in vivo* has a negligible effect on canonical miRNAs but brings an order of magnitude more siRNAs and several mirtrons (a non-canonical miRNA class) across organs (Buccheri et al., 2024). Our secondary RNAi model was Dicer^O^ expressed from a transgene inserted into the ROSA26 locus (Taborska, 2024). Although the second model was not working as originally designed, it delivered higher Dicer^O^ expression and increased RNAi in multiple organs (Taborska, 2024).

**Figure 1.**
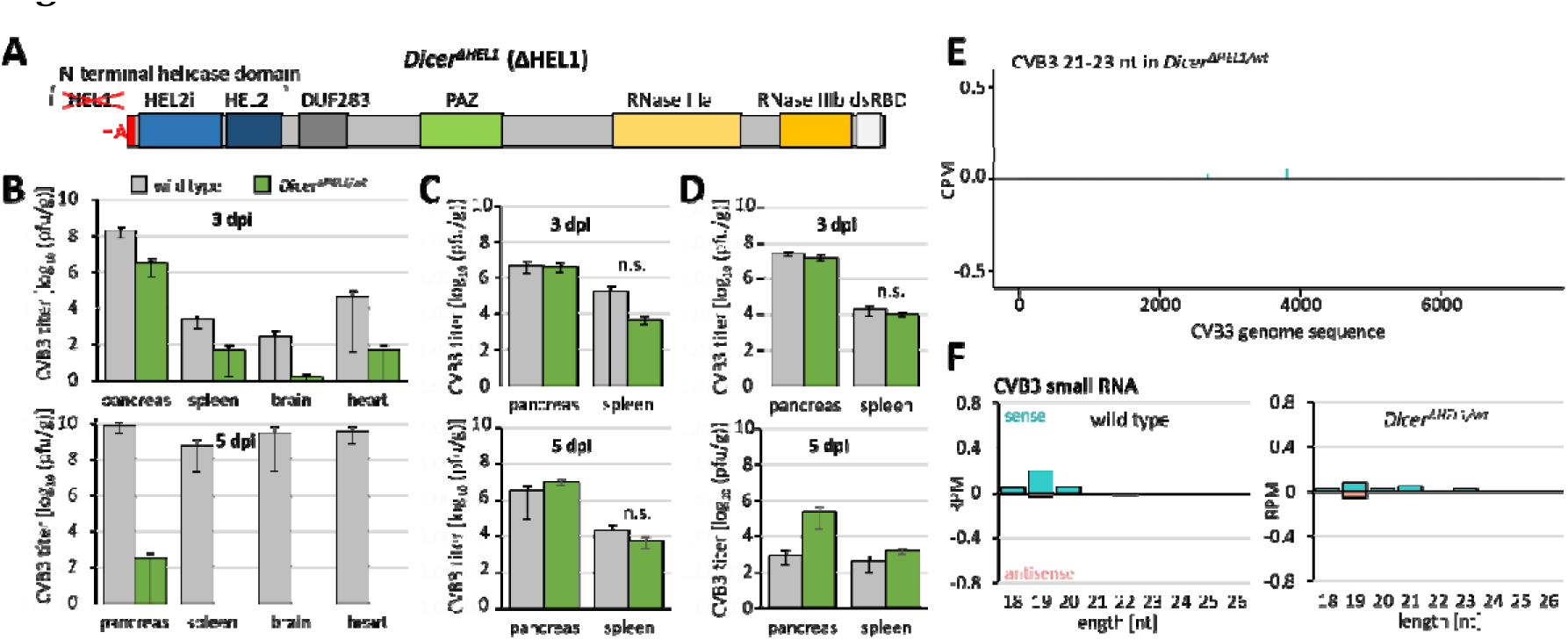
Infection of *Dicer*^Δ*HEL1/wt*^ with CVB3 (*Picornaviridae*). (A) Schematic depiction of Dicer^ΔHEL1^ protein isoform expressed from a modified endogenous *Dicer* locus. (B-E) Viral titers at 3 dpi and 5 dpi in different infection experiments: (B) Viral titers at 3 dpi and 5 dpi in the pilot experiment, in which 4 weeks old wild type mice developed systemic infection at 5dpi but *Dicer*^Δ*HEL1/wt*^ mice did not; n = 5 animals per genotype/timepoint. (C) Combined results of the next two infection experiments attempting to reproduce data from B. 7 and 12 animals were used at 3 dpi for wild type and *Dicer*^Δ*HEL1/wt*^ mice, respectively, 5 animals per genotype were used at 5 dpi. n.s. = statistically not significant reduction of viral titers in *Dicer*^Δ*HEL1/wt*^ mice (one-tailed t-test). (D) Viral titers at 3 dpi and 5 dpi after infection of 3 weeks old mice (n = 5 animals per genotype/timepoint). (E) A coverage plot for CVB3 21-23 nt small RNAs derived from + and -strand (displayed above and below the central line, respectively). The x-axis represents the viral genome sequence, the y-axis represents counts per million (CPM) of 18-32 nt reads from two different infected spleens 3 dpi from four weeks-old mice analyzed in the panel C. (F) Distribution of frequencies of small RNAs of different lengths mapping onto the CVB3 genome in infected spleens. Error bars = SD. RPM = reads per million of 18-32 nt reads.

For infection, four RNA viruses, which have been researched as mammalian pathogens in mice, were available to us. There were two viruses from the family *Picornaviridae*: CVB3 (strain Nancy) and ECMV, positive single-stranded RNA viruses lacking an envelope. Their targeting by RNAi was examined in cell culture previously. CVB3 was neither processed by full-length Dicer into vsiRNAs nor targeted by RNAi in AGO2-dependent manner in cultured cells (Schuster et al., 2019; Schuster et al., 2017) while EMCV was reported to be targeted by RNAi in ESCs (Maillard et al., 2013). The next one was the tick-borne encephalitis virus (TBEV), a positive single stranded enveloped virus from the family *Flaviviridae*, which is targeted by RNAi in the tick host (Schnettler et al., 2014). The last virus was the lymphocytic choriomeningitis virus (LCMV) from the family *Arenaviridae*, a family of segmented negative single-stranded RNA enveloped viruses, whose targeting by endogenous mammalian RNAi was not studied but was sensitive to targeting by exogenous siRNAs (Sanchez et al., 2005).

## Results

### Absence of antiviral effects in *Dicer*^Δ*HEL1/wt*^ mice

We set out to investigate whether a physiologically expressed Dicer^ΔHEL1^ variant supports innate antiviral immunity. To this end, we have exposed *Dicer*^Δ*HEL1/wt*^ mice to four different viruses (picornaviruses CVB3 and ECMV, a flavivirus TBEV, and an arenavirus LCMV) and investigated whether these viruses were recognized and targeted by enhanced RNAi in *Dicer*^Δ*HEL1/wt*^ mice.

### CVB3 (strain Nancy)

The first tested virus was the CVB3 (strain Nancy (Lindberg et al., 1987)), an ssRNA (+) virus, which belongs to the genus *Enterovirus* of the family *Picornaviridae* (Garmaroudi et al., 2015). In collaboration with the Enterovirus laboratory at the Slovak Medical University infections of *Dicer*^Δ*HEL1/wt*^ mice (CD1 outbred background) were performed. In the first experiment, four weeks old mice (5 mice per genotype and timepoint) were infected with 0.5 ml of 10^5^ TCID_50_/ml by intraperitoneal injection and organs were collected at 3 and 5 dpi (Fig. 1B-D). We observed significant reduction of the viral titer in *Dicer*^Δ*HEL1/wt*^ animals (Fig. 1B). This finding was corroborated by histology and RT-qPCR analysis of viral RNA in infected mice, where mild to severe infiltration was observed in the acinar cells of the pancreatic tissue as compared to the uninfected controls where the infiltration was absent (Fig. S1A). However, reduction in viral copies and replicating viruses was not reproduced in any subsequent infection of four weeks old (Fig. C) and three weeks old mice (Fig. D).

Furthermore, we did not detect virus-derived 21-23 nt siRNAs (vsiRNAs) in infected spleen (viral titer ∼10^4^ plaques/g) of juvenile mice. Putative vsiRNAs were not detectable at the sequencing depth of 20 million reads (Fig. 1E, F and S1B).

### EMCV

The second tested virus was the picornavirus EMCV, which has tropism for the heart (reviewed in (Carocci and Bakkali-Kassimi, 2012)) and its targeting by RNAi was reported in ESCs (Maillard et al., 2013). EMCV was a strong candidate for testing antiviral RNAi because efficient RNAi and endo-siRNA production was observed in hearts of *Dicer*^Δ*HEL1/wt*^ mice (Buccheri et al., 2024). In collaboration with the Laboratory of Arbovirology from the Biology Centre of the Czech Academy of Sciences, we performed a pilot experiment with three C57Bl/6 young adult mice per group injected subcutaneously with 10^3^ PFUs and tested progression of the EMCV infection at 2 dpi and 3 dpi. However, analysis of RNA from infected hearts showed several fold increased levels of viral RNA in *Dicer*^Δ*HEL1/wt*^ mice relative to wild type siblings (Fig. 2A). Small RNA-seq of infected hearts at 3 dpi showed low amounts of 21-23 nt reads mostly localized to the 5’ end of the viral genome sequence (Fig. 2B), reminiscent of siRNA production from blunt-end substrates in mammalian cells (Demeter et al., 2019) and consistent with EMCV siRNA distribution along the viral sequence from infected ESCs (Maillard et al., 2013). Processing of EMCV into vsiRNAs was supported by a minor peak of 22 nt RNA species in RNA-seq data in wild type and *Dicer*^Δ*HEL1/wt*^ mice (Fig. 2C) and phasing analysis showing positive signal for sense 21-23 nt RNA in the 22 nt register and a two nucleotide shift for antisense 21-23 nt RNA, which corresponds to published results from infected ESCs (Maillard et al., 2013) (Fig. 2D).

**Figure 2.**
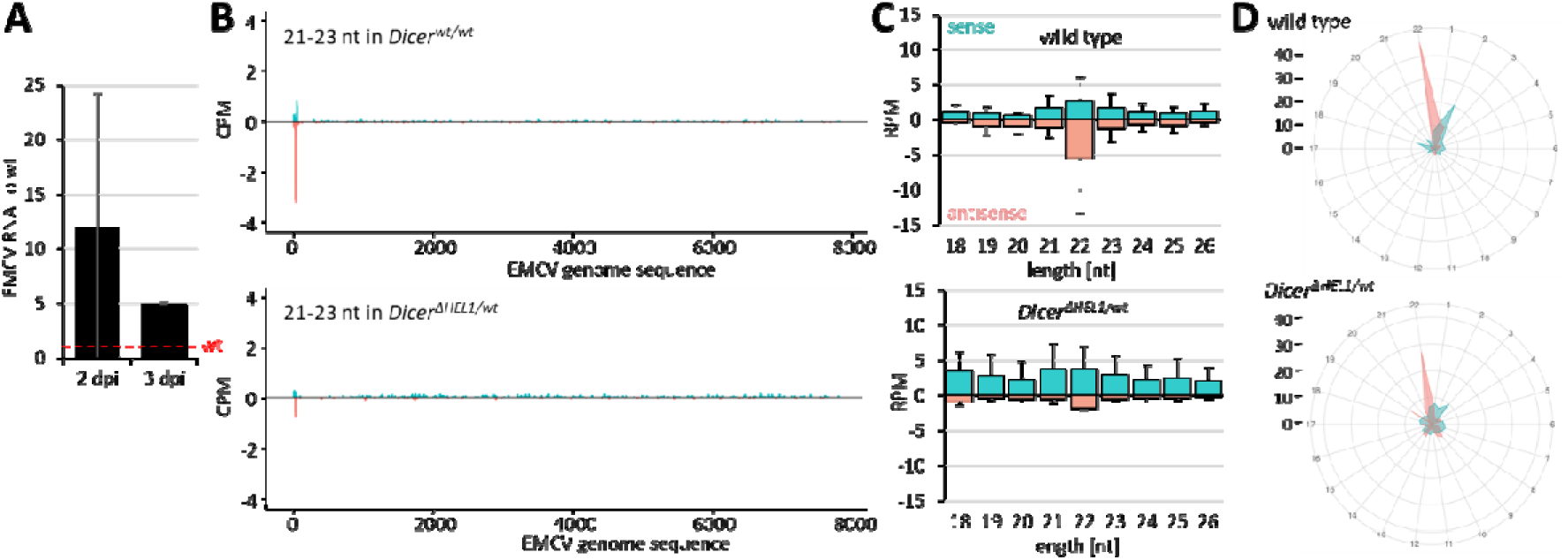
Infection of *Dicer*^Δ*HEL1/wt*^ with EMCV (*Picornaviridae*). (A) EMCV viral RNA in heart of infected *Dicer*^Δ*HEL1/wt*^ 6 weeks-old mice is several folds higher than in wild type littermates. Animals were infected subcutaneously with 10^3^ PFUs. (2 dpi: 3 animals per genotype; 3 dpi: 2 animals per genotype). Error bars = SD for 2dpi, range of two values for 3dpi. (B) A coverage plot for ECMV 21-23 nt small RNAs represents combined data for two different infected hearts 3 dpi for sense (+) and antisense (-) small RNAs mapped to the viral genome sequence. (C) Distribution of frequencies of small RNAs of different lengths mapping onto the EMCV genome in infected hearts (n=2, 3 dpi). Error bars = range of values. (D) Phasing analysis of 21-23 nt RNAs mapping onto the EMCV genome sequence. The viral genome sequence was divided into of 22 possible phased registers in both, sense and antisense orientation, and 21-23 nt reads mapping to the sequence were counted in these registers based on the position of their 5’ nucleotide. Abundance of reads in each register is displayed as distance from the center, which indicates read percentage within each register. The 5′ first EMCV nucleotide defines the register no. 1. Radar plots show 21-23 nt reads assigned to 22 possible registers along the entire EMCV sense (iris blue) and antisense (salmon) strands in wild type and *Dicer*^Δ*HEL1/wt*^ infected hearts.

However, abundance of putative EMCV vsiRNAs was very low, which contrasted with three orders of magnitude higher abundance vsiRNAs observed in infected ESCs (Maillard et al., 2013). Furthermore, there was no apparently increased abundance of EMCV-derived 21-23 nt RNAs between *Dicer*^Δ*HEL1/wt*^ and wild type mice (Fig. 2C). This contrasted with efficient siRNA production from expressed dsRNA in *Dicer*^Δ*HEL1/wt*^ mice (Buccheri et al., 2024) and suggested poor accessibility/processing of the viral dsRNA by Dicer *in vivo*. Furthermore, increased Dicer activity *in vivo* in the heart apparently promoted the viral infection as indicated by the increased amount of EMCV RNA in heart. The cause of the pro-viral effect is unclear but appears miRNA-independent as there were minimal miRNome changes in *Dicer*^Δ*HEL1/wt*^ hearts of infected animals (Fig. S1D), which concerned primarily low-expressed mirtrons and miRNA passenger strands. In any cause, because of the pro-viral effect, we did not investigate the EMCV infection model further.

### TBEV

The third tested virus was TBEV. In collaboration with the Laboratory of Arbovirology, we conducted two experiments with young adult C57Bl/6 mice (6-7 weeks old) infected by subcutaneous injection of TBEV (10^3^ PFUs). *Dicer*^Δ*HEL1/wt*^ mice lost weight and developed symptoms comparably to wild type animals (Fig. 3A,B). There was no difference in the TBEV titer between *Dicer*^Δ*HEL1/wt*^ and wild type controls at 10 dpi (Fig. 3C). RNA-seq analysis of the brain at 10 dpi provided little evidence for TBEV vsiRNAs. TBEV-derived 21-23 nt RNAs were mostly coming from the sense strand (Fig. 3D and S2A). However, size distribution of RNA fragments did not show enrichment of 21-23 nt RNAs in comparison to longer and shorter small RNAs in wild type or *Dicer*^Δ*HEL1/wt*^ infected brains (Fig. 3E), suggesting that small RNAs from the sense strand are mostly degradation fragments. Notably, analysis of antisense small RNAs, which were two orders of magnitude less abundant than sense fragments, revealed a distinct peak of 21-23 nt RNAs derived from the minus strand in *Dicer*^Δ*HEL1/wt*^ brains (Fig. 3F). This peak was not observed in TBEV-infected wild type brains (Fig. 3F). This implied that RNAi machinery in brain of *Dicer*^Δ*HEL1/wt*^ mice cleaves TBEV dsRNA but that does not significantly affect viral replication. Consistent with that, 21-23 nt vsiRNAs in *Dicer*^Δ*HEL1/wt*^ mice brains appeared too low-abundant for efficient RNAi (Buccheri et al., 2024) when considering abundance of all TBEV RNA fragments.

**Figure 3.**
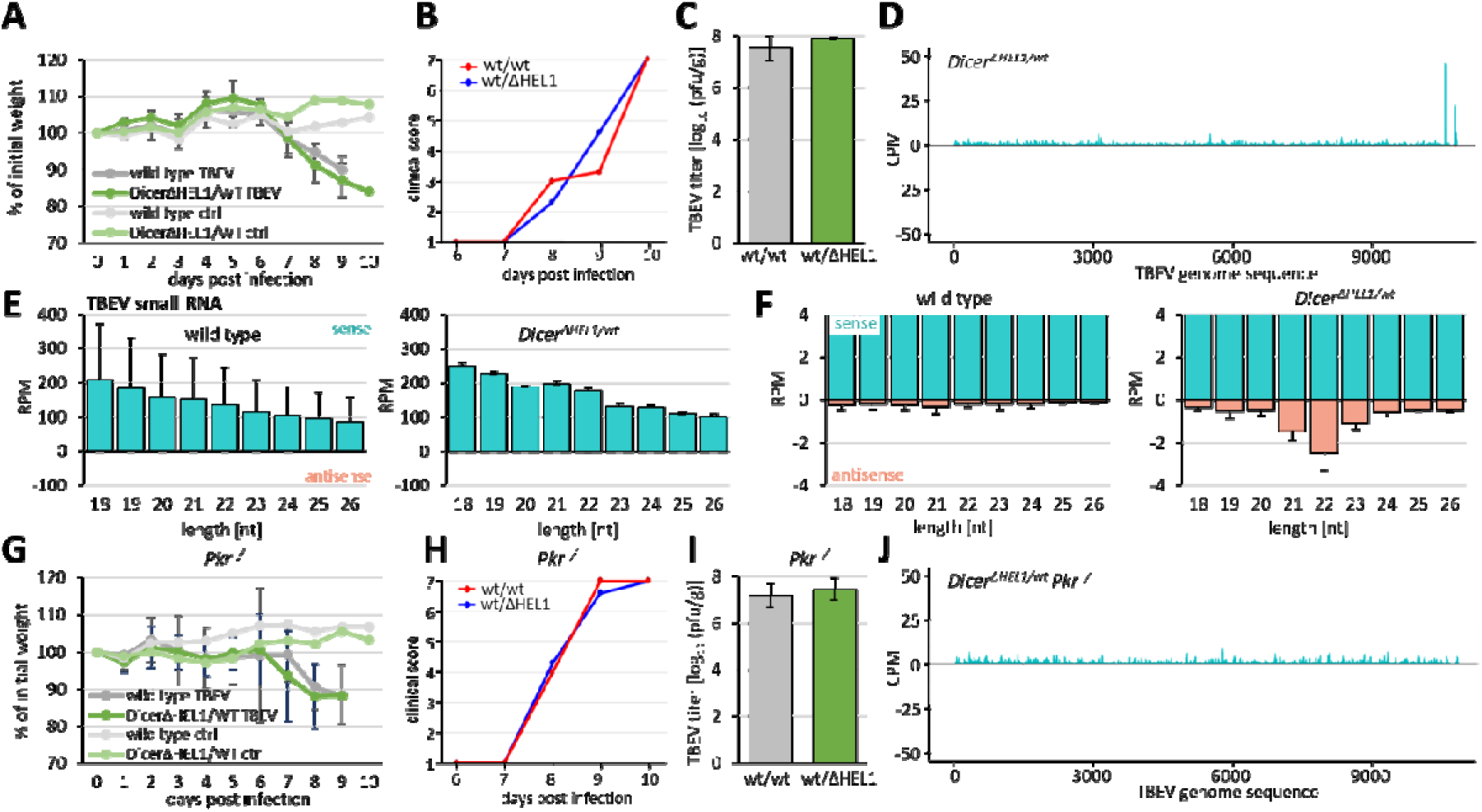
Infection of *Dicer*^Δ*HEL1/wt*^ (A-F) and *Dicer*^Δ*HEL1/wt*^ *Pkr^−/−^* (G-J) and wild type mice with TBEV (*Flaviviridae*). Animals were infected subcutaneously with 10^3^ PFU. (A) and (G) Loss of weight of infected animals. n= 3. Error bars = SEM. (B) and (H) Development of seve rity of infection. Clinical score 1 = healthy, 2 = piloerection, 3 = hunched back, 4 = paralysis (one leg), 5 = paralysis (two legs), 6 = moribund, 7 = dead. (C) and (I) TBEV viral titers obtained from infected brains from animals of the specified genotypes at 10 dpi (7 weeks-old, n = 3 animals per genotype/timepoint). Error bars = SD (D) and (J) Coverage plots for TBEV 21-23 nt small RNAs in brains of *Dicer*^Δ*HEL1/wt*^ and *Dicer*^Δ*HEL1/wt*^ *Pkr^−/−^* mice at 10 dpi. The panel (D) combines data from three different infected brains, panel (J) shows data from a single brain. (E) Distribution of frequencies of small RNAs of different lengths mapping onto the TBEV genome in infected brains (n = 3; 10 dpi). Error bars = SD. (F) The same data as in the panel E but shown at a scale where 21-23 nt antisense RNAs are visible. Antisense 21-23 nt RNAs in *Dicer*^Δ*HEL1/wt*^ are significantly more abundant than in wild type brains (two-tailed t-test p-value <0.05).

We next tested how the absence of the protein kinase R (PKR, official symbol EIF2AK2), a key innate immunity dsRNA sensor, would impact virus targeting by RNAi *in vivo*. The rationale for this experiment stemmed from the fact that experiments in cell culture (ES and 3T3 cells) reported that the loss of PKR stimulated siRNA production and RNAi (Demeter et al., 2019; Kennedy et al., 2015). We infected wild type and *Dicer*^Δ*HEL1/wt*^ mice, which were also homozygous for deletion of dsRNA binding-domain encoding exons of *Pkr*. *Pkr* mutants showed a similar loss of weight as mice with normal *Pkr* (Fig. 3G vs. 3A), but died one day earlier (Fig. 3H). *Pkr* mutants reached the same TBEV viral titers in the post-mortem brain as their *Pkr* wild type counterparts (Fig. 3I vs. 3C) and there was no significant difference in the viral titer in the post mortem brain between wild type and *Dicer*^Δ*HEL1/wt*^ in the *Pkr* mutant background (Fig. 3I). Small RNA-seq of 3, 6, and 9 dpi samples did not support TBEV targeting by RNAi as 21-23 nt TBEV-derived RNAs had the above-mentioned strong asymmetry towards the (+) strand and did not show clear enrichment of 21-23 nt small RNA species (Fig. 3J and S2B).

It was reported that TBEV, flaviviruses related to TBEV, and tick-borne Langat virus (LGTV) produced virus-derived 22 nt small RNAs in tick cells (Schnettler et al., 2014). Indeed, we have observed higher abundance of specific small RNAs from the 3’ end of the viral genomic sequence, which was increased in *Dicer*^Δ*HEL1/wt*^ samples (Fig. S2A). It has been also reported that tick borne flaviviruses may suppress RNAi through a protein (Qiu et al., 2020) or a structured RNA (Schnettler et al., 2014). However, despite this, *Dicer*^Δ*HEL1/wt*^ mutants still showed significantly increased vsiRNA production (Fig. 3F). Furthermore, analysis of miRNA expression in *Dicer*^Δ*HEL1/wt*^ mutants showed highly selective miRNA changes, which did not support general inhibition of Dicer function in the brain (Fig. S2C). In fact, increased mirtron levels in brains of infected *Dicer*^Δ*HEL1/wt*^ mice but not in wild type animals document increased activity of Dicer^ΔHEL1^ (Fig. S2C).

### LCMV

The final *in vivo* tested virus was LCMV, a segmented (-)ssRNA virus from the family *Arenaviridae* with tropism for secondary lymphoid organs (reviewed in (Laposova et al., 2013)). This work was done in collaboration with the Laboratory of Adaptive Immunity from the Institute of Molecular Genetics of the Czech Academy of Sciences. The virus was delivered by intraperitoneal injection and its levels in spleen were not significantly reduced in neither *Dicer*^Δ*HEL1/wt*^ nor in *Dicer*^Δ*HEL1/wt*^ *Pkr^−/−^* mutants (Fig. 4A). Similarly to the EMCV infection, we observed 21-23 nt small RNAs generated from sense and antisense strands, particularly at the termini of viral genomic RNAs (Fig. 4B). However, small RNA analysis showed no specific enrichment of 21-23 nt small RNAs among 18-32 nt small RNAs and there was similar abundance of putative vsiRNAs in wild type animals and *Dicer*^Δ*HEL1/wt*^ mutants (Fig. 4C and Fig. S3A). Small RNA size distribution had a distinct 19 nt peak, which corresponded to small RNAs mapping to complementary termini of S and L segments (Fig. S3B) and likely represent degradation fragments with increased stability. Loss of PKR did not have any positive effect on increased enrichment of 21-23 nt small RNAs (Fig. S3C) Phasing analysis of 21-23 nt small RNAs showed a weak signal in the register 1 for both, sense and antisense 21-23 nt RNAs in the whole sequence. This signal became much stronger when the analysis was restricted only to the terminal 100 nucleotides. This implied that a minor fraction of 21-23 nt reads, particularly from the termini, might be produced by Dicer. At the same time, enhanced RNAi in *Dicer*^Δ*HEL1/wt*^ was insufficient to mediate an antiviral effect. Thus, to gain more insights into the effect of RNAi on LCMV, we turned to ESCs where we could investigate how Dicer activity may limit antiviral effects.

**Figure 4.**
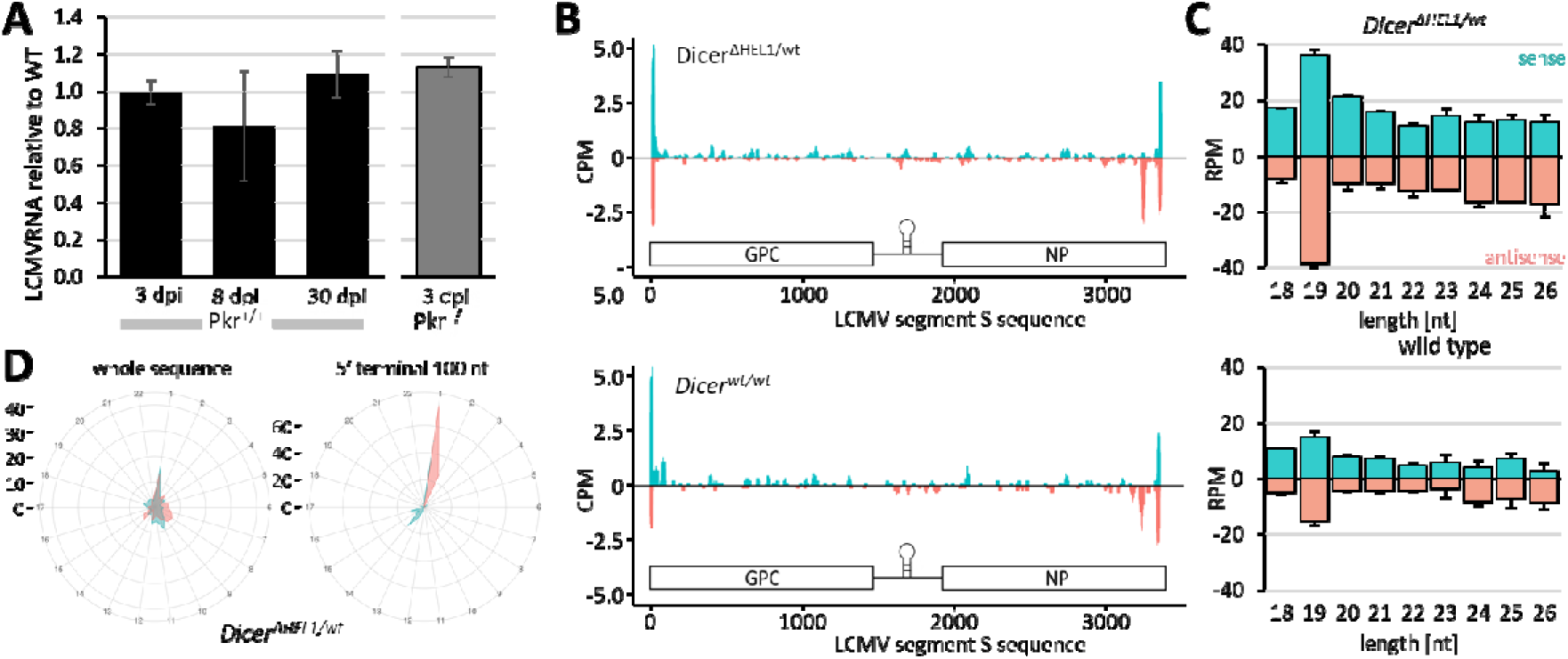
Infection of *Dicer*^Δ*HEL1/wt*^ with LCMV (*Arenaviridae*). (A) LCMV RNA in spleens from 11-17 weeks-old *Dicer*^Δ*HEL1/wt*^ and *Dicer*^Δ*HEL1/wt*^ *Pkr*^-/-^ animals infected with LCMV at 3 dpi, 8 dpi or 30 dpi relative to WT (n = 3 animals per genotype/timepoint). Animals were injected with 2x10^5^ PFUs except of the 30 dpi timepoint where animals were injected with 10^6^ PFUs of LCMV C13 intravenously. Error bars = SEM. (B) Coverage plots for 21-23 nt small RNAs mapping to the S segment of the LCMV genome. Each plot depicts combined data obtained from two different spleens at 3 dpi. (C) Analysis of distribution of lengths of small RNAs derived from the segment S at 3dpi in spleen of infected *Dicer*^Δ*HEL1/wt*^ and *Dicer^wt/wt^* (n = 2; error bars = range of values). (D) Phasing analysis of 21-23 nt RNAs mapping onto LCMV sequence. The left diagram is based on analysis of all mapped 21-23 nt RNAs while the right one took into the account only those mapping to 100 nucleotides at the 5′ terminus of the genomic RNA.

### Higher levels of truncated Dicer variants target LCMV in ESCs

We employed two ESC lines with high Dicer activity produced in the lab previously: *Dicer*^Δ*HEL1/*Δ*HEL1*^ (Buccheri et al., 2024; Zapletal et al., 2022) and *Dicer^O-3^* ESC lines (Flemr et al., 2013). Unlike mice, ESCs withstand *Dicer*^Δ*HEL1/*Δ*HEL1*^ homozygosity. The *Dicer^O-3^* line carries a stable expression of the oocyte-specific Dicer^O^ variant in the absence of expression of endogenous Dicer (Flemr et al., 2013) and is expressing even more truncated Dicer than *Dicer*^Δ*HEL1/*Δ*HEL1*^ ESCs (Fig. 5A).

**Figure 5.**
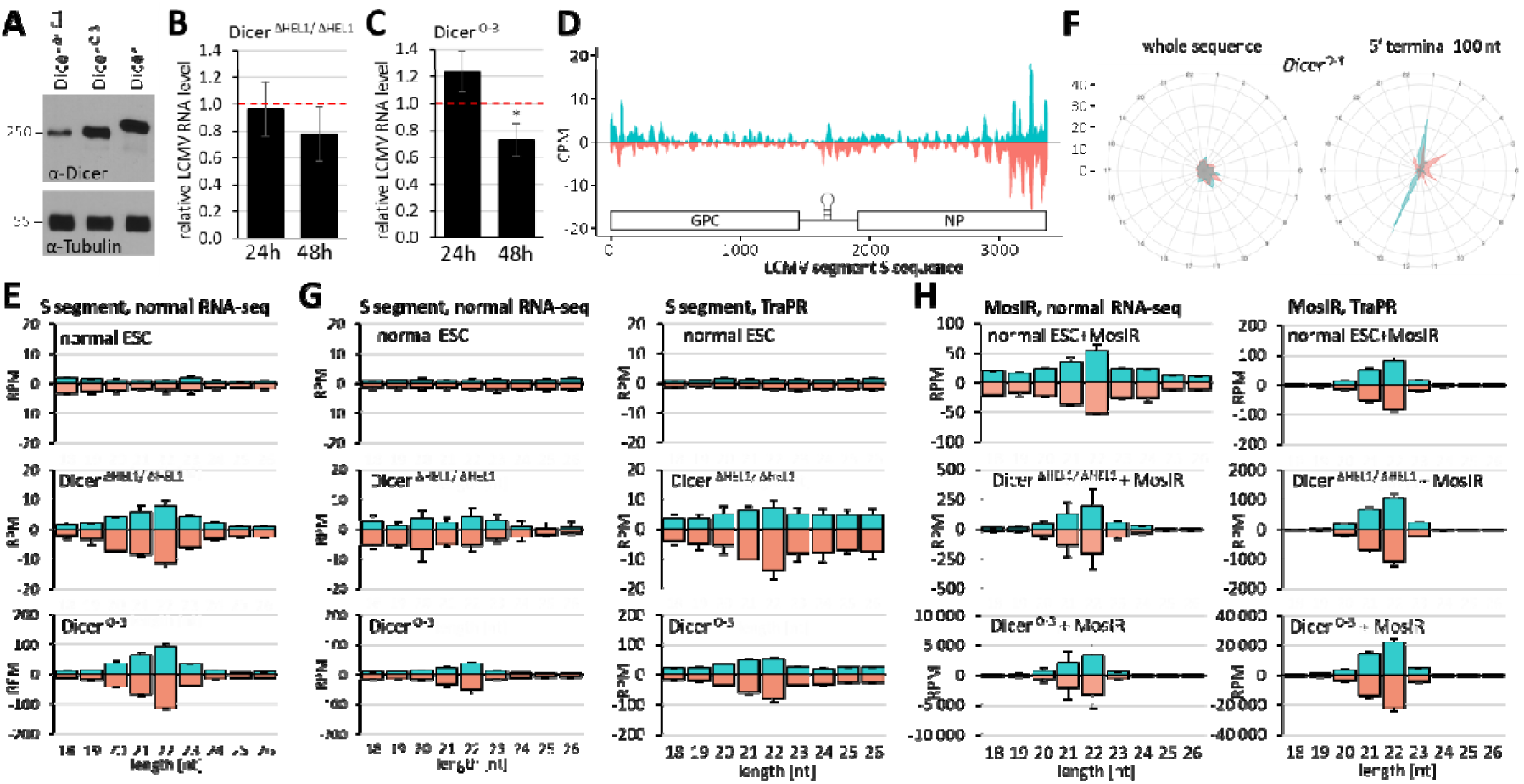
Analysis of ESCs lines expressing different levels of Dicer^ΔHEL^ shows RNAi-dependent antiviral effects. (A) Western blot analysis of Dicer protein levels in Dicer^ΔHEL1^ homozygote, Dicer^O-3^ ESC line and in unmodified ESCs (Dicer). (B) LCMV RNA levels 24 and 48 hpi (MOI = 0.01) in *Dicer*^Δ*HEL1/*Δ*HEL1*^ ESCs relative to the infected parental ESC line. Erros bars = SEM. (C) LCMV RNA levels 24 and 48 hpi (MOI 0.01) in Dicer^O-3^ ESCs line levels relative to Dicer^S-4^ ESC line expressing comparable levels of full-length Dicer (Flemr et al., 2013). Erros bars = SEM. * p-value <0.05 (D) Coverage plot for 21-23 nt small RNAs mapping to the S segment of the LCMV genome depicts data from infected Dicer^O-3^ ESCs at 24 hpi (n=2). (E) Small RNA length distribution analysis from parental WT ESCs, *Dicer*^Δ*HEL1/*Δ*HEL1*^ and *Dicer^O-3^* (n = 2; 24 hpi). (F) Phasing analysis of 21-23 nt RNAs from infected *Dicer^O-3^* ESCs mapping onto LCMV sequence. The left diagram is based on analysis of all mapped 21-23 nt RNAs while the right one took into the account only those mapping to 100 nucleotides at the 5′ terminus of the genomic RNA. (G-H) Analysis of AGO-bound LCMV siRNAs. RNA sequencing was performed in parallel from total RNA isolated from infected cells 24 hpi (MOI = 0.1, n=2) either directly (left panel) or with TrapR methodology to isolate AGO-associated small RNAs (right panel). (H) Small RNA length distribution from parental WT, *Dicer*^Δ*HEL1/*Δ*HEL1*^ and *Dicer^O-3^* ESCs transfected with a plasmid encoding the MosIR dsRNA hairpin (24 h post transfection, n=2). Before sequencing, AGO-associated small RNAs were isolated with TrapR methodology (right panel). Error bars in panels E, G, H= range of values.

Infection with MOI 0.01 of ESC lines with increased Dicer activity showed minor but statistically significant reduction of LCMV RNA in infected cells: 22.1% in *Dicer*^Δ*HEL1/*Δ*HEL1*^ ESCs (Fig. 5B) and 26.9% in Dicer^O-3^ ESC line (Fig. 5C). While the general profile of virus-derived 21-23 nt small RNAs also showed a higher signal around termini of the viral genome (Fig. 5D), there was clear enrichment of LCMV-derived 21-23 nt RNAs in *Dicer*^Δ*HEL1/*Δ*HEL1*^ ESCs (Fig. S4A) and even higher abundance of LCMV-derived 21-23 nt RNAs in *Dicer^O-3^* ESC relative to control ESCs (Fig. 5E and S4B). However, LCMV substrate was clearly a limiting factor for siRNA production as well because infection of *Dicer*^Δ*HEL1/*Δ*HEL1*^ ESCs with MOI 1.0 yielded even higher even higher abundance of LCMV-derived 21-23 nt (Fig. S4C).

Similarly to the analysis of LCMV-derived 21-23 nt RNAs from infected mice (Fig. 4F), phasing analysis of *Dicer^O-3^* ESCs showed a strong signal in the register 1 for both, sense and antisense 21-23 nt RNAs when the analysis was restricted to the terminal 100 nucleotides (Fig. 5F). Signal on the sense strand in the register 1 suggests that the genomic RNA at 5’ end might be a nucleotide shorter than the annotated sequence. Interestingly, a second signal on the sense strands came from the register 13 (Fig. 5F). We hypothesize that this signal comes from a sense strand, which was cleaved by antisense siRNA but still replicated by the viral RNA polymerase, thus creating a blunt end dsRNA starting at this position. Notably, when the whole S segment sequence was used, phasing analysis of 21-23 nt RNAs from in *Dicer^O-3^*ESCs did not show a dominant signal in any register (Fig. 5F). At the same time, RNA-seq from *Dicer^O-3^* ESCs showed abundant 21-23 nt LCMV-derived RNA pool (Fig. 5E and S4B) and notably enhanced decoration of the S segment by sense and antisense 21-23 nt RNAs in the coverage plot (Fig. 5D and S4B). These data suggest that in addition to phased 21-23 nt RNAs produced its termini, abundant highly active Dicer may initiate LCMV dsRNA processing by stochastic endonucleolytic cleavage, thus generating a pool of vsiRNAs, which are not matching a specific register.

To test whether the observed 21-23 nt RNAs are bound by Argonaute proteins, we compared normal RNA-seq with RNA-seq of RNA isolated with TraPR isolation method, which purifies RNAs associated with the RISC effector complex (Grentzinger et al., 2020). As a positive control for siRNA production, we included ESCs transfected with a MosIR plasmid, which is expressing long dsRNA hairpin activating RNAi (Flemr et al., 2013; Svoboda et al., 2001). After sequencing, we observed clear peaks of 21-23 nt small RNAs in TraPR-isolations from *Dicer*^Δ*HEL1/*Δ*HEL1*^ and *Dicer^O-3^* ESCs but not *Dicer^wt/wt^*, which were having approximately two times higher reads per million (RPM) counts than normal RNA-seq, suggesting that they were indeed bona fide LCMV-derived vsiRNAs (Fig. 5G). Consistent with higher expression of a truncated Dicer variant, *Dicer^O-3^* ESCs yielded about an order of magnitude more vsiRNAs (Fig. 5G). At the same time, MosIR 21-23 nt RNAs from TraPR isolation were much more abundant (Fig. 5H). MosIR siRNAs were detectable in normal ESCs at abundance similar to that of LCMV siRNAs in *Dicer^O-3^* ESCs. In comparison to normal ESCs, *Dicer*^Δ*HEL1/*Δ*HEL1*^ and *Dicer^O-3^* ESCs exhibited one and two orders of magnitude higher MosIR siRNA levels, respectively. Taken together, ESC infections with LCMV provided evidence that increased abundance of a Dicer variant lacking the HEL1 domain induces vsiRNA production and repression of LCMV.

### Higher Dicer**^Δ^**^HEL1^ expression is antiviral *in vivo*

To test whether LCMV could be affected *in vivo* by increasing Dicer activity beyond that achieved in *Dicer*^Δ*HEL1/wt*^, we took advantage of a transgene Tg(EGFP-lox66-pCAG-lox71i-Dicer^O-HA^-T2A-mCherry), for simplicity referred to as *Dicer^Tg(O-HA)^* hereafter (Fig. 6A). This transgene was produced for Cre-inducible expression of C-terminally HA-tagged Dicer^O^ (Dicer^O-HA^) from the CAG promoter (Taborska, 2024). However, analysis of the transgenic mouse line revealed that the uninduced allele had leaky expression, which varied across organs (Fig. 6B) and differed from previously reported expression of CAG (Miyazaki et al., 1989; Okabe et al., 1997). The leakage was the highest in testes where it was observed in meiotic and postmeiotic cells (Taborska, 2024). Analysis of total Dicer mRNA expression in different organs of animals carrying one or two copies of the transgene suggested 1-3 times increased *Dicer* transcript level, except of the testis, where the leakage was much stronger (Fig. 6C). These data suggested that *Dicer^Tg(O-HA)^* mice homozygous for the transgene express higher truncated Dicer level than *Dicer*^Δ*HEL1/wt*^ mice and we thus examined how they would respond to LCMV infection.

**Figure 6.**
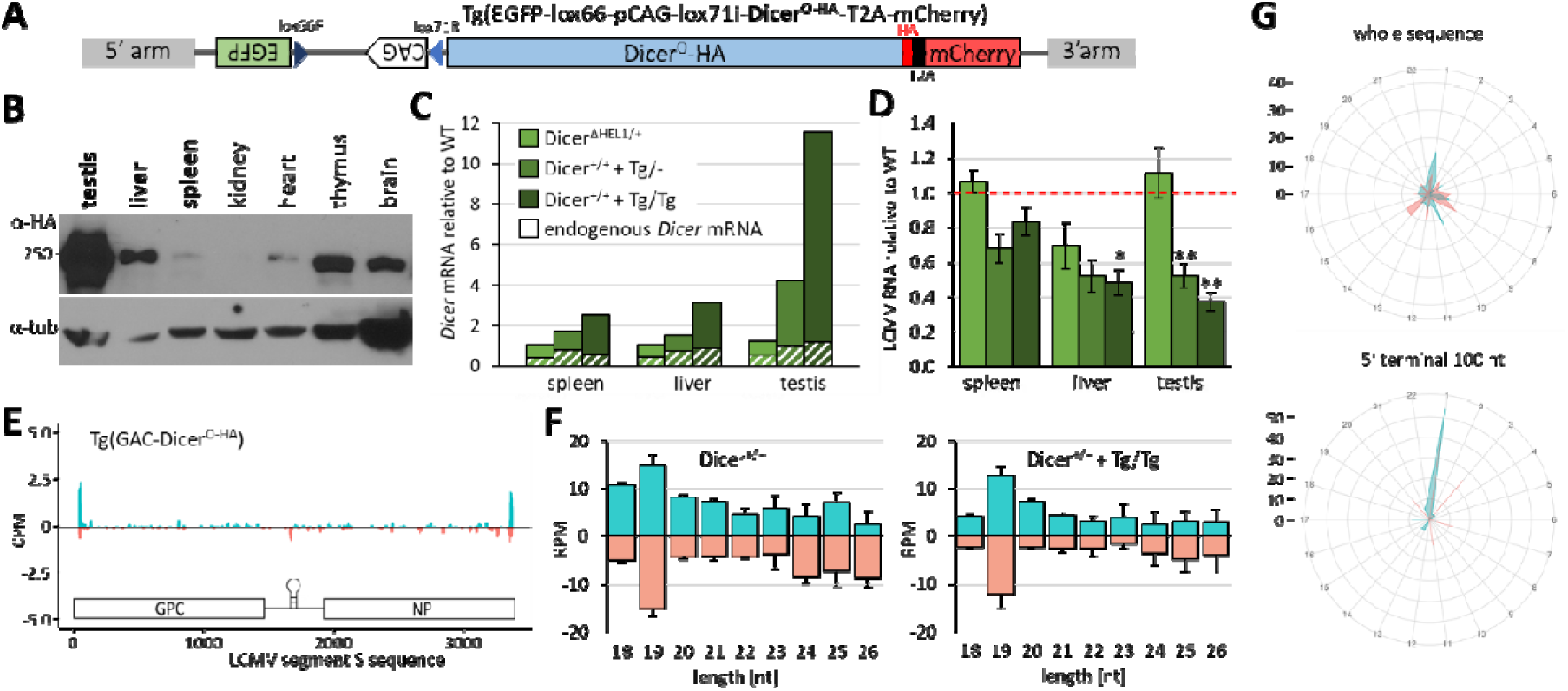
Antiviral effects *in vivo* in *Dicer^Tg(O-HA)^* transgenic mice overexpressing truncated Dicer. (A) A scheme of the inducible Dicer transgene. (B) Leaky transgene expression in *Dicer^Tg(O-HA)^* mice. Despite the CAG promoter is oriented antisense to the *Dicer* coding sequence, the transgene has leaky Dicer expression, which varies among organs. The expression differs from the CAG promoter expression pattern observed in other transgenes(Buccheri et al., 2024; Nejepinska et al., 2012; Okabe et al., 1997). The western blot was adopted from (Taborska, 2024) for readers convenience. (C) Estimation of total and endogenous *Dicer* mRNA levels in spleen, liver and testis of mice with the three indicated genotypes. RNA from organs of one animal was analyzed by RT-qPCR in a technical triplicate, shown is the median value. The white-hatched parts in the bars correspond to the estimated level of the endogenous full-length Dicer mRNA amplified by a different set of primers. (D) LCMV RNA in the indicated organs from 11-12 weeks-old infected *Dicer*^Δ*HEL1/wt*^, *Dicer^+/+^+ ^Tg/-^* and *Dicer^+/+^+ ^Tg/Tg^* animals relative to wild type littermates. Data are from two independent experiments 3 animals per genotype per experiment were injected with 2.5x10^6^ PFU of LCMV Arm intraperitoneally, organs were collected at 3 dpi. Error bars = SEM. * and ** indicate p-value <0.05 and <0.01 from a two-tailed t-test, respectively. (E) Coverage plot for 21-23 nt small RNAs mapping to the S segment of the LCMV genome depicts combined data obtained from two different spleens at 3 dpi. Animals were infected with 2.5x10^6^ PFU of LCMV Arm intraperitoneally. (F) Analysis of distribution of lengths of small RNAs derived from viral RNA in spleen of infected *Dicer*^Δ*HEL1/wt*^*, Dicer^+/+^+ ^Tg/Tg^* and WT littermates (n = 2; error bar = range of values). (G) Phasing analysis of 21-23 nt RNAs from infected *Dicer^+/+^ +^Tg/Tg^* spleen mapping onto the LCMV sequence. The upper diagram is based on analysis of all mapped 21-23 nt RNAs while the right one took into the account only those mapping to 100 nucleotides at RNA termini.

Remarkably, when we infected mice carrying one or two copies of the *Dicer^Tg(O-HA)^* transgene, we have observed 20-30% reduction of LCMV RNA levels in spleen and 50-60%, reduction in liver and testes and (Fig. 6D). The difference was statistically significant in transgenic homozygotes in the liver and testes and in transgenic hemizygotes in testes (Fig. 6C). This is a solid yet relatively weak antiviral effect considering that strong antiviral effects reduce viruses by orders of magnitude. Notably, this antiviral effect was not accompanied with a marked increase in 21-23 nt RNA levels in spleen *Dicer^Tg(O-HA)^* homozygous mice relative to *Dicer^wt/wt^* mice (Fig. 6E and F). In fact, 22 nt LCMV RNAs seemed to be less abundant than most of the other sizes in infected spleens (Fig. 6E and F). LCMV 21-23 nt RNA in testes and liver were below 1 RPM and considered too low for analysis (data not shown). Phasing analysis revealed the same picture as infection of *Dicer*^Δ*HEL1/wt*^, i.e. signal in the register 1, which became much stronger when the analysis was restricted to the 5′ terminal 100 nucleotides (Fig. 6G vs. 4D). This would imply that a large portion of 21-23 nt LCMV RNAs in spleen come from the termini and it would be consistent with the coverage plot (Fig. 6E).

## Discussion

Our results provide a new framework for antiviral RNAi and bring several important implications. We have genetically engineered the *Dicer*^Δ*HEL1*^ allele in mice to establish an *in vivo* model with enhanced RNAi, which could fill a gap in the research on the antiviral potential of mammalian RNAi. The gap stemmed from the fact that Dicer and AGO2 are essential for the miRNA pathway, so it is impossible to study antiviral RNAi *in vivo* in their absence; this includes the loss of catalytic activity of AGO2 (Bernstein et al., 2003; Cheloufi et al., 2010; Liu et al., 2004). The *Dicer*^Δ*HEL1/wt*^ mouse offers an experimental alternative where heterozygosity leaves the canonical miRNA pathway essentially intact while the mouse exhibits an order of magnitude higher siRNA production and more efficient RNAi ubiquitously (Buccheri et al., 2024). We analyzed four viruses whose selection primarily stemmed from availability in collaborating laboratories and established procedures for infecting mice *in vivo*. The four viruses were used for testing whether the enhanced RNAi in *Dicer*^Δ*HEL1/wt*^ mutants would be sufficient to provide detectable antiviral effects. In fact, it was previously reported that EMCV was targeted by RNAi in ESCs (Maillard et al., 2013), TBEV is targeted by RNAi in the tick host (Schnettler et al., 2014) and that other *Picornaviridae* and *Flaviviridae* viruses are targeted by RNAi in mammals (Fang et al., 2021; Kakumani et al., 2013; Qiu et al., 2017; Qiu et al., 2020; Xu et al., 2019; Zhang et al., 2022). However, *Dicer*^Δ*HEL1/wt*^ mice did not show any reproducible enhanced defense against the tested viruses. We observed suppression of CVB3 in the first experiment, which was not reproduced in subsequent three independent infections. We cannot exclude that the genetic background could play a role as shown before (Srivastava et al., 2009). In contrast to EMCV, TBEV and LCMV experiments done in an inbred C57Bl/6 background, CVB3 experiments were performed using CD1 outbred genetic background onto which was initially crossed the *Dicer*^Δ*HEL1/wt*^ allele, which was prepared in R1 ESCs, which come from 129 background. In the very first CVB3 experiment, wild type littermates developed systemic infection while *Dicer*^Δ*HEL1/wt*^ mice did not (Fig. 1B). In subsequent experiments, systemic infection did not develop in any animals. We speculate that the CD1 background with residual 129 background could have affected the outcome. In any case, this initial CVB3 result is an example of experimental variability, which could not be reproduced.

RNAi requires efficient processing of long dsRNA into siRNA, which is not provided by the mammalian full-length Dicer. The N-terminal helicase is the key structural element of Dicer, which facilitates miRNA biogenesis and prevents efficient long dsRNA conversion into siRNA (Flemr et al., 2013; Chakravarthy et al., 2010; Ma et al., 2008; Zhang et al., 2002).

Accordingly, Dicer variants with a modified N-terminal helicase have been examined in mammalian antiviral RNAi studies. Among them, stands out the so-called N1 variant lacking HEL1 and HEL2i domains (Baldaccini et al., 2024; Kennedy et al., 2015; Montavon et al., 2021). The N1 variant has increased siRNA biogenesis while it does not interact with dsRBP proteins that modulate Dicer function (Montavon et al., 2021; Wilson et al., 2015). It was also shown in HEK293 cells that the N1 Dicer has antiviral activity against SINV, SFV and EV71 (but not against VSV or SARS-CoV-2), which involves RNAi-independent stimulation of interferon and inflammatory response pathways (Baldaccini et al., 2024). Another reported endogenous Dicer isoform is the antiviral Dicer (AviD) isoform. It was identified by PCR and lacks the HEL2i domain through skipping exons 7 and 8 during splicing (Poirier et al., 2021). AviD was shown to support antiviral RNAi in HEK293 cells against SINV and ZIKV and it was proposed to provide antiviral protection in intestinal stem cells (Poirier et al., 2021). The physiological role of AviD is unknown but it is apparently non-essential in healthy animals as suggested by phenotype analysis of *Dicer^SOM/SOM^* mice, which lack AviD and could be used for testing how significant could be the antiviral role of AviD *in vivo* (Buccheri et al., 2024; Taborska et al., 2019).

The Dicer^ΔHEL1^ protein variant used in our study is an HA-tagged functional equivalent of Dicer^O^ variant. The Dicer^ΔHEL1^ isoform was structurally and functionally thoroughly investigated (Buccheri et al., 2024; Demeter et al., 2019; Zapletal et al., 2022). It lacks the HEL1 subdomain, which causes higher siRNA production, but retains HEL2i, which is important for protein-protein interaction (Liu et al., 2018). Since the *Dicer*^Δ*HEL1*^ mutation preserved endogenous transcription control of Dicer, *Dicer*^Δ*HEL1/wt*^ mice represent the minimal possible intervention to stimulate siRNA production *in vivo*. However, we observed little if any vsiRNA production and no antiviral effects for CVB3, EMCV, TBEV, and LCMV in *Dicer*^Δ*HEL1/wt*^ mice.

Based on observed putative siRNA levels, we hypothesized that a single *Dicer*^Δ*HEL1*^ allele may not be sufficient to produce Dicer activity necessary for antiviral effects. To test if further increased Dicer activity could have any measurable effect, we further examined LCMV infections in *Dicer*^Δ*HEL1/*Δ*HEL1*^ ESCs and, since *Dicer*^Δ*HEL1/*Δ*HEL1*^ animals are not viable, we produced a transgenic model with higher Dicer activity where Dicer^O^ was expressed on top of the normal endogenous Dicer expression. Indeed, we detected antiviral effects in both cases supporting the idea that antiviral RNAi has higher Dicer activity threshold than can be delivered by a single *Dicer*^Δ*HEL1*^ allele *in vivo* but this threshold can be reached with ectopic expression of a truncated Dicer as was the case of Dicer^O-3^ ESC line and Dicer^Tg(O-HA)^ transgene in LCMV infections.

Apart from Dicer, amount and accessibility of viral long dsRNA are likely significant limitations for generating enough vsiRNAs for efficient RNAi. vsiRNA RPM values in *Dicer*^Δ*HEL1/*Δ*HEL1*^ and *Dicer^O-3^* ESCs were relatively low when compared to high siRNA levels from expressed long dsRNA (Fig. 5G and H) and siRNA levels correlating with RNAi silencing in cultured cells or *in vivo* (Buccheri et al., 2024; Demeter et al., 2019; Flemr et al., 2013). However, canonical RNAi represents a steady-state system of independently expressed long dsRNA and its complementary target, which is degraded via RISC and thus requires high amount of AGO-loaded siRNAs. Antiviral RNAi operates on a replicating system where even a small repression would have a leverage effect over replication cycles and repression may involve both: cleavage of a replicating virus by Dicer and targeting viral transcripts by AGO-bound vsiRNAs. Yet, it is still puzzling that a 21-23 nt peak in RNA-seq of LCMV infected samples was observed in ESCs but not *in vivo*, despite we have observed reduced amount of LCMV RNA in *Dicer^Tg(O-HA)^* mice. This contrasted with ESC experiments where vsiRNAs were readily visible as a 21-23 nt peak in the *Dicer*^Δ*HEL1/*Δ*HEL1*^ line (Fig. 5E). Several factors may contribute to this discrepancy. First, the antiviral effect may occur without accumulating vsiRNAs as it may also come from the Dicer-mediated cleavage of viral RNA and not just from the AGO2-mediated cleavage required for canonical RNAi.

Furthermore, target-mediated decay (Ameres et al., 2010; De Madrid and Porterfield, 1969) might promote stronger depletion of AGO2-bound vsiRNAs in organs. Another contributing factor could be that ESC samples have reduced non-specific RNA fragments because RNAs from released viral particles and cellular debris are removed when cells are washed with PBS before harvesting while a whole infected organ is directly used for RNA isolation.

Taken together, our research on mice expressing Dicer^ΔHEL1^ variants does not contradict antiviral RNAi in mammals but reframes it and highlights several important considerations for future research. It is important to recognize that mammalian endogenous RNAi is not absent but mostly ineffective because of low Dicer activity, limiting dsRNA substrate levels, and activity of sequence-independent dsRNA responses.

Conversely, providing high Dicer activity, accessible dsRNA substrate, and reduction of sequence-independent dsRNA responses facilitate RNAi. It is thus possible to find experimental conditions to show RNAi functionality, especially in experimental systems tolerating high Dicer expression, controlling dsRNA abundance and with reduced interference from sequence-independent dsRNA responses. RNAi may be a negligible nuisance for most viruses but some may be sensitive to it. Future research should further delineate the actual physiological potential of endogenous antiviral RNAi *in vivo* and utility of transient activation of RNAi via a truncated Dicer variant to provide an additional layer of innate immunity.

## Materials and Methods Biological resources *Animals*

Animal experiments were carried out in accordance with the Czech law and were approved by the Institutional Animal Use and Care Committee (approval no. 34-2014, 29-2019; and 86-2020).

*Dicer*^Δ*HEL1*^ mice were produced as previously described (Zapletal et al., 2022). For viral infection experiments were used viable heterozygotes, which were characterized in detail previously (Buccheri et al., 2024). *Pkr (Eif2ak2)* mutant mice and *Tg(PFGE-GAC-Dicer^O-HA^-mCHERRY)* transgenic mice on C57Bl/6 background were described elsewhere (Buccheri et al., 2024). Animals were genotyped by PCR. Briefly, tail biopsies were lysed in DEP-25 DNA Extraction buffer (Top-Bio) according to manufacturer’s instructions. 1 µl aliquots were mixed with primers and Combi PPP Master Mix (Top-Bio) for genotyping PCR. Genotyping primers are provided in Table S1. For analyses, organs collected from sacrificed animals were either directly used for analysis or stored at -80° C for later use.

### Cell lines

Mouse ESCs were cultured in 2i-LIF media: KO-DMEM (Gibco) supplemented with 15% fetal calf serum (Sigma), 1x L-Glutamine (Thermo Fisher Scientific), 1x non-essential amino acids (Thermo Fisher Scientific), 50 µM β-Mercaptoethanol (Gibco), 1000 U/mL LIF (Isokine), 1 µM PD0325901, 3 µM CHIR99021(Selleck Chemicals), penicillin (100 U/mL), and streptomycin (100 µg/mL) at 37°C in a humidified atmosphere with 5% CO_2_.

Human lung adenocarcinoma A549 cells (ECACC 86012804) were cultured in Dulbecco’s Modified Eagle’s Medium (DMEM) containing 10% fetal bovine serum (FBS; Biosera), 1% penicillin, 1% streptomycin, and 1% glutamine (Biowest) at 37°C in a humidified atmosphere with 5% CO_2_.

Hamster BHK-21 and mouse 3T3 cells were cultured in Dulbecco’s Modified Eagle’s Medium (DMEM) containing 10% fetal bovine serum (FBS; Biosera), 1% penicillin, 1% streptomycin at 37°C in a humidified atmosphere with 5% CO_2_.

### Viral infections

#### CVB3

*In vivo* were conducted on the original CD1 genetic background of the *Dicer*^Δ*HEL1/wt*^ mice in collaboration with the Enterovirus laboratory at the Slovak Medical University. The virus was propagated in Vero cells (origin Public Health Institute, Bratislava) and 0.5 ml of 10^5^ TCID_50_/1ml virus was injected intraperitoneally, per mouse. Mice were sacrificed and samples collected 3 and 5 days post infection (dpi); in two experiments organs were also collected 7 dpi, 11 dpi and 45 dpi.

#### EMCV

The EMCV isolate BCCO_50_0517 (GenBank: PP841942.1) was passaged on VERO/E6 prior to infection. This virus was provided by the Collection of Arboviruses, Biology Centre of the Czech Academy of Sciences (https://arboviruscollection.bcco.cz). *Dicer*^Δ*HEL1/wt*^ mice on the C57Bl/6 background were infected subcutaneously into the scruff of the neck with 10^3^ PFUs. Mice were euthanized by cervical dislocation and heart samples were collected 2 and 3 dpi. The samples (2-5mm^3^ of the organ) were placed into 600 ul Trizol and frozen (-80°C) for later RNA isolation.

#### TBEV

The TBEV strain Hypr (GenBank MT228627.1) was passaged five times in the brains of suckling mice before it was used in this study. This strain was provided by the Collection of Arboviruses, Biology Center of the Czech Academy of Sciences (https://arboviruscollection.bcco.cz). *Dicer*^Δ*HEL1/wt*^ and *Dicer*^Δ*HEL1/wt*^ *Pkr^−/−^* mice on the C57Bl/6 background were infected subcutaneously into the scruff of the neck with 10^3^ PFUs. Mice were observed daily for body weight and clinical score (healthy, piloerection, hunched back, one leg paralyzed, two legs paralyzed, moribund, dead) as previously (Agudelo et al., 2021) until 10 dpi. Mice were euthanized by cervical dislocation and brain samples were collected and homogenized as previously (Pokorna Formanova et al., 2019). Brains were weighed individually and then homogenized in sterile PBS (1:1) using the TissueLyser II (Qiagen). 100 ul of the brain suspension was transferred to 600 ul Trizol and frozen (-80°C) for subsequent RNA isolation. The remaining homogenate was clarified by centrifugation at 14,000×g for 10 min at 4 °C and then used for virus titration using a plaque assay on A549 cells.

#### LCMV

Viral stocks of LCMV, strains Armstrong (Arm) and Clone 13 (C13), were propagated in BHK-21 cells as described previously (Horkova et al., 2023). These strains were obtained from Prof. Daniel Pinschewer (University Hospital Basel, Switzerland). The viral titer in aliquots was determined by LCMV Focus Forming Assay. Briefly, 3T3 cells were infected with different dilutions of the virus supernatant. Viral antigens were detected with rat anti-LCMV nucleoprotein antibody (Clone VL-4; BioXCell, Cat. N.: BE0106, Lot: 787521S1, diluted 1:500), and visualized by a color reaction using secondary goat anti-rat IgG HRP antibody (diluted 1:500) 48 hours post infection(hpi). For LCMV acute infection, each mouse was injected intraperitoneally 2x10^5^ or 2.5x10^6^ PFUs of LCMV Arm. At 3 dpi or 8 dpi, mice were euthanized by cervical dislocation and organ samples were collected for total RNA isolation. All used animals had the C57Bl/6 background. For LCMV chronic infection, each mouse was injected intravenously 10^6^ PFUs of LCMV C13. At 30 dpi, mice were euthanized by cervical dislocation and organ samples were collected for total RNA isolation. Mouse ESCs were infected with Arm at an multiplicity of infection (MOI) of 0.01 or 1.0 and collected 24 or 48 hpi.

### Tissue histology

After formalin fixation, tissues were dehydrated using graded alcohols, cleared with xylene, and infiltrated with paraffin wax. Appropriate small tissue pieces were collected in 4% formaldehyde and embedded in paraffin wax cassettes. The tissues from paraffin-embedded blocks were cut into 4-7µm thick slices on a microtome and mounted from warm distilled water (40°C) onto microscope silane coated slides (Super Frost Plus, Menzel-Glaser).

Sections were allowed to dry overnight at 40°C. Prior to staining, tissue sections were deparaffinized in xylene and rehydrated by stepwise washes in decreasing ethanol/H_2_O ratio (96%, 70%) for 5 min in each and in distilled water. The slides were stained with hematoxylin solution (Mayer’s solution) for 10 min washed in running water, then stained with Eosin, the slides were treated with 70% ethanol for 20 sec, 90% ethanol for 20 sec, 100% ethanol for 1 min and xylene for 3 min dried, mounted with xylene-based mounting media and coverslips (Wang et al., 2017).

### CVB3 virus titration of the organs suspension

To determine the replicating CVB3 titers in organs of infected mice Hep-2 cells were used. Cells were grown monolayers in 96-well flat bottom plates in MEM(E) supplemented with 1% HEPES, antibiotics and 10% fetal bovine serum.

Serial 10-fold dilutions of the organ suspensions or of the serum samples with MEM(E) (supplemented with 1% HEPES, antibiotics and 2% fetal bovine serum) were made in 96-well U bottom plates (8 wells for each dilution) and 100 μl of diluted suspensions were transferred to the grown monolayers (the medium from the grown monolayers was first discarded). The plates were incubated in a CO_2_ incubator at 37°C and checked on days 4-5 under the light microscope. Titers were expressed as 50% tissue culture infectious dose (TCID_50_) following TCID_50_ values, calculated according to Karbeŕs method (Hierholzer and Killington, 1996).

### TBEV Plaque assay

To determine TBEV titers, A549 cells were used following a modified version of a previously described protocol (De Madrid and Porterfield, 1969). Tenfold dilutions of the infectious samples were placed in 24-well plates and incubated with A549 cell suspension (1.2 x 10^5^ cell per well) for 4 hours at 37 °C and 5% CO2. The samples were then covered with overlay mixture (1.5% carboxymethylcellulose in complete culture media). After 5 days, the plates were washed with PBS and stained with naphthalene black. Virus-produced plaques were counted, and the titers were expressed as PFUs/ml.

### Transfection

For transfection, cells were plated on a 24-well plate, grown to 50 % density and transfected using Lipofectamine 3000 (Thermo Fisher Scientific) according to the manufacturer’s protocol.

### Western blot

Mouse organs and ES cells were homogenized mechanically in RIPA lysis buffer supplemented with 2x protease inhibitor cocktail set (Millipore) and loaded with SDS dye. Protein concentration was measured by Bradford assay (Bio-Rad) and 80 μg of total protein was used per lane. Proteins were separated on 5.5% polyacrylamide (PAA) gel and transferred on PVDF membrane (Millipore) using semi-dry blotting for 60 min, 35 V. The membrane was blocked in 5% skim milk in TBS-T, Dicer was detected using rabbit polyclonal anti-Dicer antibody #349 (Sinkkonen et al., 2010) (a gift from Witold Filipowicz, dilution 1:5000), anti-HA rabbit primary antibody (Cell Signaling, #3724, dilution 1:1,000) and incubated overnight at 4°C. Secondary anti-Rabbit-HRP antibody (Santa-Cruz #sc-2357, dilution 1:50,000) was incubated 1 h at room temperature. For TUBA4A detection, samples were separated on 10% PAA gel and incubated overnight at 4 °C with anti-Tubulin (Sigma, #T6074, dilution 1:10,000). HRP-conjugated anti-mouse IgG binding protein (Santa-Cruz, #sc-525409, dilution 1:50,000) was used for detection. Signal was developed on films (X-ray film Blue, Cole-Parmer #21700-03) using SuperSignal West Femto Chemiluminescent Substrate (Thermo Scientific).

### RNA isolation

Infected cells, non-infected cells and mouse organs were washed with PBS, homogenized in Qiazol lysis reagent (Qiagen) and total RNA was isolated by Qiazol-chloroform extraction and ethanol precipitation method (Toni et al., 2018).

### RT-qPCR analysis

For Dicer expression analysis, aliquots of 3 µg of total RNA were treated by Turbo DNA-*free^TM^* kit (Invitrogen) according to the manufacturer’s instructions. Next, aliquots of 1 µg of total DNAse-treated RNA were used for cDNA synthesis by LunaScript RT SuperMix Kit (New England Biolabs) according to the manufacturer’s instructions. For LCMV and EMCV RNA quantification, cDNA synthesis was performed directly after RNA isolation. A 1 µl cDNA aliquot and the Maxima SYBR Green qPCR master mix (Thermo Fisher Scientific) were used for the qPCR reaction. qPCR was performed on LightCycler 480 (Roche) in technical triplicates for each biological sample. Average Ct values of the technical replicates were normalized using the ΔΔCt method to three housekeeping genes *Hprt*, *Alas*, and *B2mg* for mouse organs and *Hprt* or *B2mg* for mouse ESCs. *eEF1a1* was used as a housekeeping gene for one 3 dpi analysis of an LCMV infection of *Dicer*^Δ*HEL1/wt*^. A list of the primers used for qPCR is provided in Supplementary Table.

### Isolation of RNA silencing complex (RISC)-associated small RNAs

ESCs infected with LCMV at a MOI of 0.1 or transfected with a plasmid expressing a MosIR dsRNA hairpin were used for isolation of RISC complexes by Trans-kingdom, rapid, affordable Purification of RISCs (TrapR, Lexogen, Austria) according to manufacturer’s instructions 24 hpi and 48 hpi, respectively. RISC-associated small RNAs were extracted by a mixture of acidic phenol, chloroform and isoamylalcohol and precipitated by cold isopropanol. RISC-associated small RNAs were immediately used for small RNA library preparation.

### Small RNA sequencing (RNA-seq)

After isolation, RNA quality was verified by electrophoresis on 1% agarose gel and RNA concentration was determined by Qubit Broad Range Assay (Invitrogen). Small RNA libraries were prepared using NEXTFLEX® Small RNA-Seq Kit v3 for Illumina (PerkinElmer) according to the manufacturer’s protocol; 3′ adapter ligation was performed overnight at 20 °C, 15 cycles were used for PCR amplification and gel purification was performed for size selection. For gel purification, libraries were separated on a 2.5% agarose gel using 1× lithium borate buffer and visualized with ethidium bromide. The 150–160 bp fraction was cut off the gel and DNA was isolated using the MinElute Gel Extraction Kit (Qiagen). Final libraries were analyzed by Agilent 2100 Bioanalyzer and sequenced by 75-nucleotide single-end reading using the Illumina NextSeq500/550 platform.

### Bioinformatic analyses

RNA-seq data (Table S2) were deposited in the Gene Expression Omnibus database under accession ID GSE273338.

### Mapping of small RNA-seq data

Small RNA-seq reads were trimmed in two rounds using fastx-toolkit version 0.0.14 (http://hannonlab.cshl.edu/fastx_toolkit) and cutadapt version 1.8.3 (Martin, 2011). First, 4 random bases were trimmed from left side:

~~~
fastx_trimmer -f 5 -i {INP}.fastq -o {TMP}.fastq
~~~

Next, NEXTflex adapters were trimmed. Additionally, the N-nucleotides on ends of reads were trimmed and reads containing more than 10% of the N-nucleotides were discarded:

~~~
cutadapt --format=“fastq” --front=”GTTCAGAGTTCTACAGTCCGACGATCNNNN” -- adapter=”NNNNTGGAATTCTCGGGTGCCAAGG” --error-rate=0.075 --times=2 --overlap=14 -- minimum-length=12 --max-n=0.1 --output=”${TRIMMED}.fastq” --trim-n --match-read-wildcards ${TMP}.fastq
~~~

Trimmed reads were mapped to the mouse (mm10) genome with following parameters:

~~~
STAR --readFilesIn ${TRIMMED}.fastq.gz --runThreadN 4 --genomeDir ${GENOME_INDEX} -- genomeLoad LoadAndRemove --readFilesCommand unpigz -c --readStrand Unstranded -- limitBAMsortRAM 20000000000 --outFileNamePrefix ${FILENAME} --outReadsUnmapped Fastx -- outSAMtype BAM SortedByCoordinate --outFilterMultimapNmax 99999 -- outFilterMismatchNoverLmax 0.1 --outFilterMatchNminOverLread 0.66 --alignSJoverhangMin 999 --alignSJDBoverhangMin 999
~~~

### Viral small RNA analyses

Small RNA-seq reads were trimmed in two rounds using bbduk.sh version 38.87 (Bushnell, 2015). First, NEXTflex adapter were trimmed from right end:

~~~
bbduk.sh -Xmx20G threads=6 in=${FILE}.fastq.gz out=${FILE}.atrim.fastq.gz literal= TGGAATTCTCGGGTGCCAAGG stats=${FILE}.atrim.stats overwrite=t ktrim=r k=21 rcomp=f mink=10 hdist=1 minoverlap=8
~~~

Next, 4 random bases from both sides of reads were trimmed:

~~~
bbduk.sh -Xmx20G threads=6 in=${FILE}.atrim.fastq.gz out=${FILE}.trimmed.fastq.gz stats=${FILE}.ftrim.stats overwrite=t forcetrimright2=4 forcetrimleft=4 minlength=18
~~~

The genome index was created by joining mouse genome .fasta file (GCA_000001635.2, mm10) with individual viral genomes (JN048469.1, NC_001479.1, NC_004291.1, NC_004294.1, U39292.1). Trimmed reads were mapped to such genome index with STAR version 2.7.10b (Dobin et al., 2013) with the following parameters:

~~~
STAR --readFilesIn ${FILE}.fastq.gz --genomeDir ${GENOME_INDEX} --runThreadN 12 -- genomeLoad LoadAndRemove --limitBAMsortRAM 20000000000 --readFilesCommand unpigz –c--outFileNamePrefix ${FILENAME} --outSAMtype BAM SortedByCoordinate -- outReadsUnmapped Fastx --outFilterMismatchNoverLmax 0.1 --outFilterMatchNmin 16 -- outFilterMatchNminOverLread 0 --outFilterScoreMinOverLread 0 –outFilterMultimapNmax 99999 --outFilterMultimapScoreRange 0 --alignIntronMax 1 –alignSJDBoverhangMin 999999999999
~~~

For viral genome coverage, only reads between 21-23 nt long were visualized. If reads were aligned to viral genomes with three or less soft-clipped nucleotides on 3’ ends of the alignment, such soft-clipped nucleotides were added to read lengths.

Mapped reads were counted using program featureCounts (Liao et al., 2014). Only reads with lengths 18-25nt were selected from the small RNA-seq data:

~~~
featureCounts -a ${ANNOTATION_FILE} -F ${FILE} -minOverlap 15 -fracOverlap 0.00 -s 1 -M -O - fraction -T 8 ${FILE}.bam
~~~

The GENCODE gene set (Frankish et al., 2019) was used for the annotation of long RNA-seq data. ThemiRBase 22.1. (Kozomara et al., 2019) set of miRNAs was used for the annotation of small RNA-seq data. Statistical significance and fold changes in gene expression were computed in R using the DESeq2 package (Love et al., 2014). Genes were considered to be significantly up- or down-regulated if their corresponding p-adjusted values were smaller than 0.05.

## Statistical analyses

For statistical testing, two-tailed t-test was used.

## Acknowledgements

We thank to Markéta Dvořáková (Biology Centre, Czech Academy of Sciences) and Darina Paprckova (IMG) for technical assistance. The main funding was provided by the Czech Science Foundation EXPRO grant 20-03950X. previous development of genetically modified Dicer mouse models was funded from the European Research Council under the European Union’s Horizon 2020 research and innovation programme (grant agreement No 647403, D-FENS). Financial support of M.I.R.K. and E.S. was in part provided by the Charles University in a form of a PhD student fellowship; this work will be in part used to fulfil requirements for a PhD degree and hence can be considered “school work”. Additional funding for work in collaborating laboratories was provided by the Czech Science Foundation grant 23-08039S (to M.P) and GA22-18046S (to O.S.) and the National Institute of Virology and Bacteriology (Programme EXCELES, ID Project No. LX22NPO5103), funded by the European Union—Next Generation EU (to D.R. and O.S.). The authors also acknowledge the following services: The Czech Centre for Phenogenomics at IMG supported by the Czech Academy of Sciences RVO 68378050 and by the project LM2018126 and LM2023036 Czech Centre for Phenogenomics provided by Ministry of Education, Youth and Sports (MEYS) of the Czech Republic; The Light Microscopy Core Facility at IMG supported by MEYS – LM2023050 and RVO 68378050-KAV-NPUI; Computational resources provided by the e-INFRA CZ project (ID:90254), supported by MEYS and by the ELIXIR-CZ project (ID:90255), a part of the international ELIXIR infrastructure.

---

**Figure S1.**
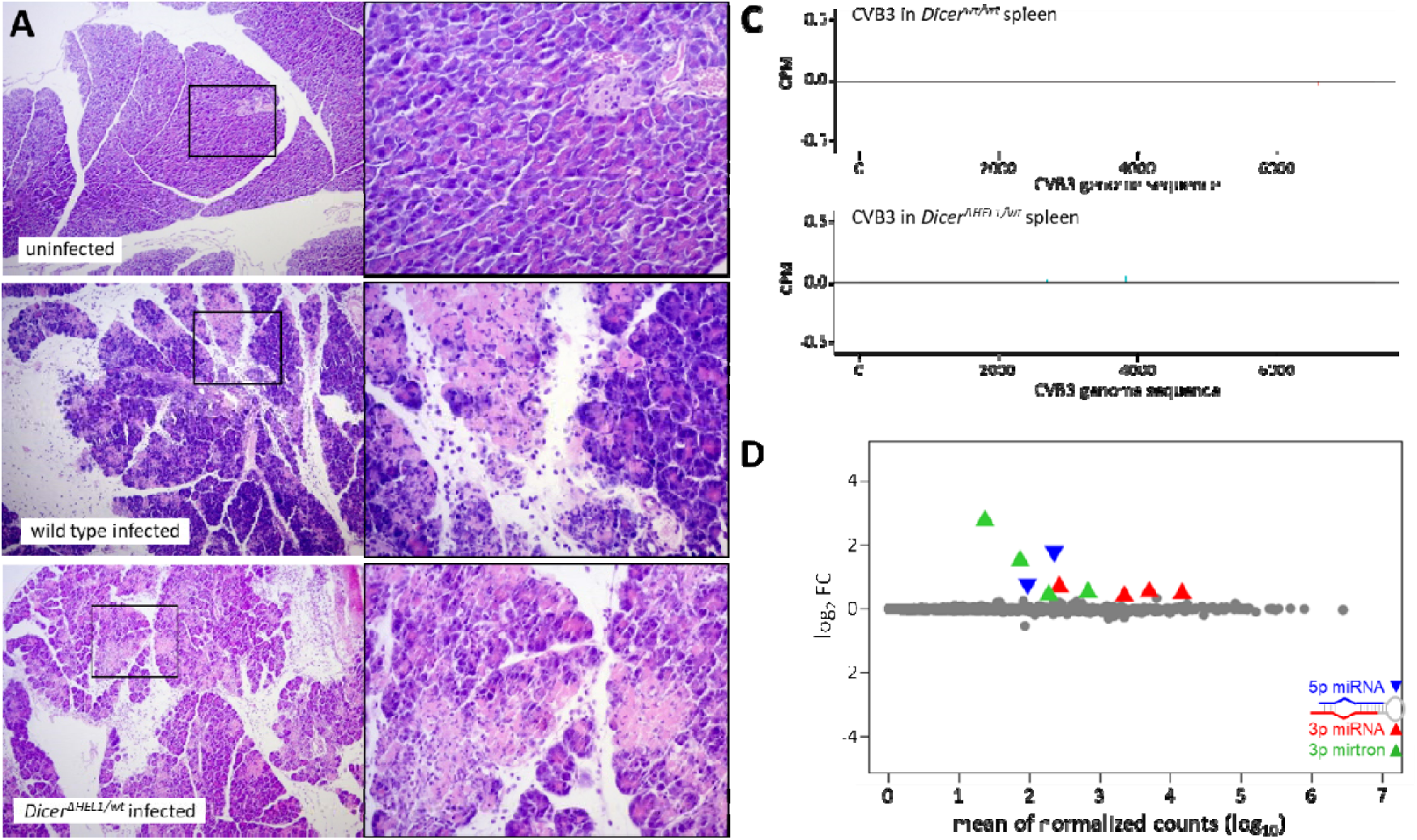
Analysis of small RNAs in mice infected with CVB3 and EMCV. (A) Histological evidence of pancreas tissue infection with CVB3 after intraperitoneal inoculation of wild type (wt) and Dicer ^Δ*HEL1/wt*^ as compared to sham infected control mice at 3dpi. Hematoxylin-Eosin (HE) staining showed control tissue without infiltration at 3 and 5 dpi, the infected wild type tissue showed infiltration and acute inflammation of the pancreatic acinar tissue at 3dpi going to chronicity at 5 dpi. (C) Coverage plot for 21-23 nt small RNAs mapping to the CVB3 genome sequence shows absence of 21-23 nt putative vsiRNA in spleen of infected mice at 3 dpi. The *Dicer*^Δ*HEL1/wt*^ panel is the same as in Fig. 1E for easier comparison with the wild type coverage plot. (D) MA plot of differentially expressed miRNAs in *Dicer*^Δ*HEL1/wt*^ EMCV infected hearts relative to wild type infected hearts, for each genotype n=3.

**Figure S2.**
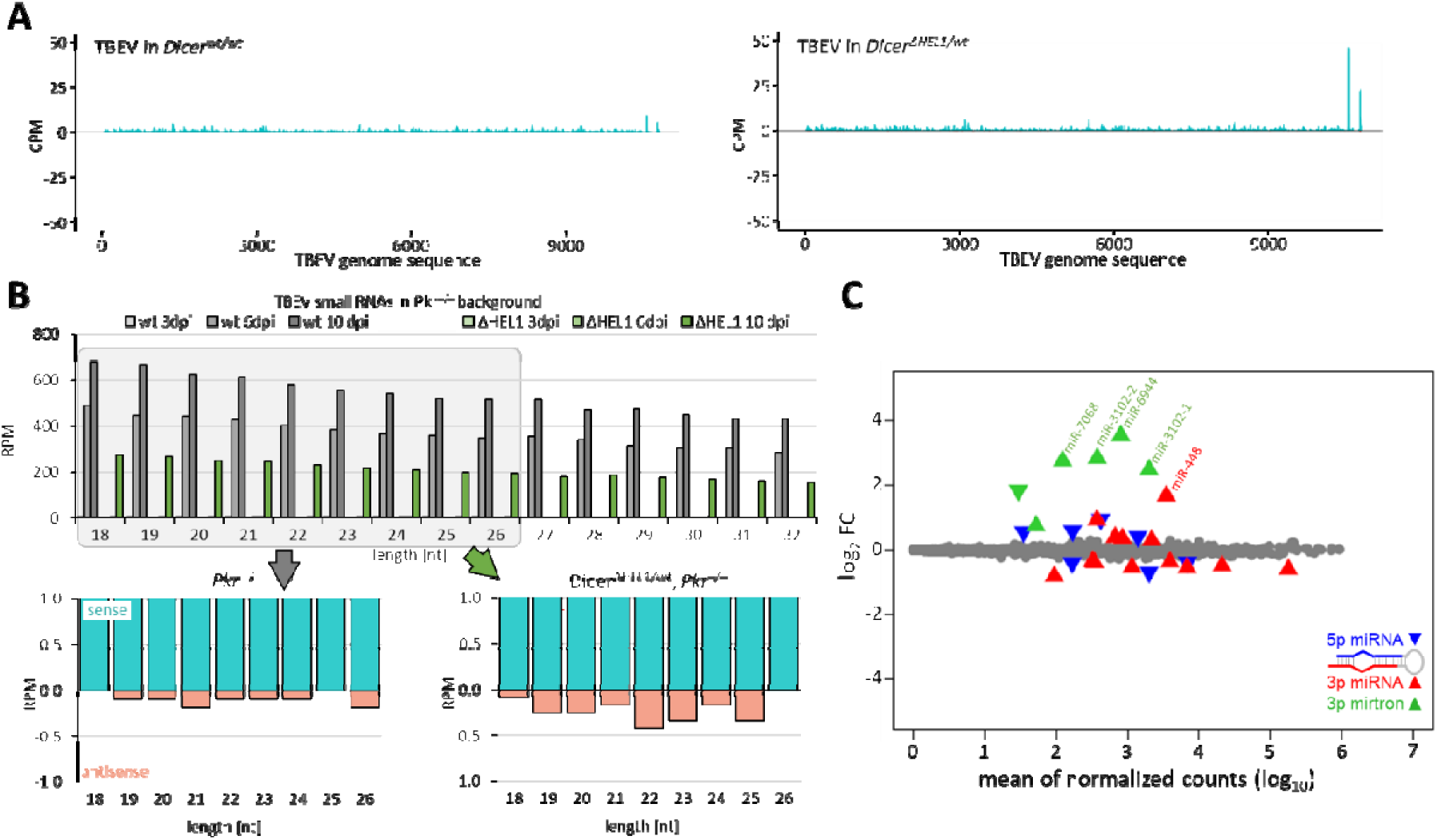
TBEV small RNA analysis. (A) Coverage plot for 21-23 nt small RNAs mapping to the TBEV genome sequence. The *Dicer*^Δ*HEL1/wt*^ panel is the same as in Fig. 3D for easier comparison with wild type samples. (B) Abundance of reads of different lengths in infected wild type and *Dicer*^Δ*HEL1/wt*^ brains lacking functional PKR. The lower graphs depict 18-26 nt distribution of sense and antisense RNAs in wild type and *Dicer*^Δ*HEL1/wt*^ brains lacking functional PKR at 10 dpi. (C) MA plot of differentially expressed miRNAs in *Dicer*^Δ*HEL1/wt*^ TBEV infected brains relative to wild type infected brains, for each genotype n=3.

**Figure S3.**
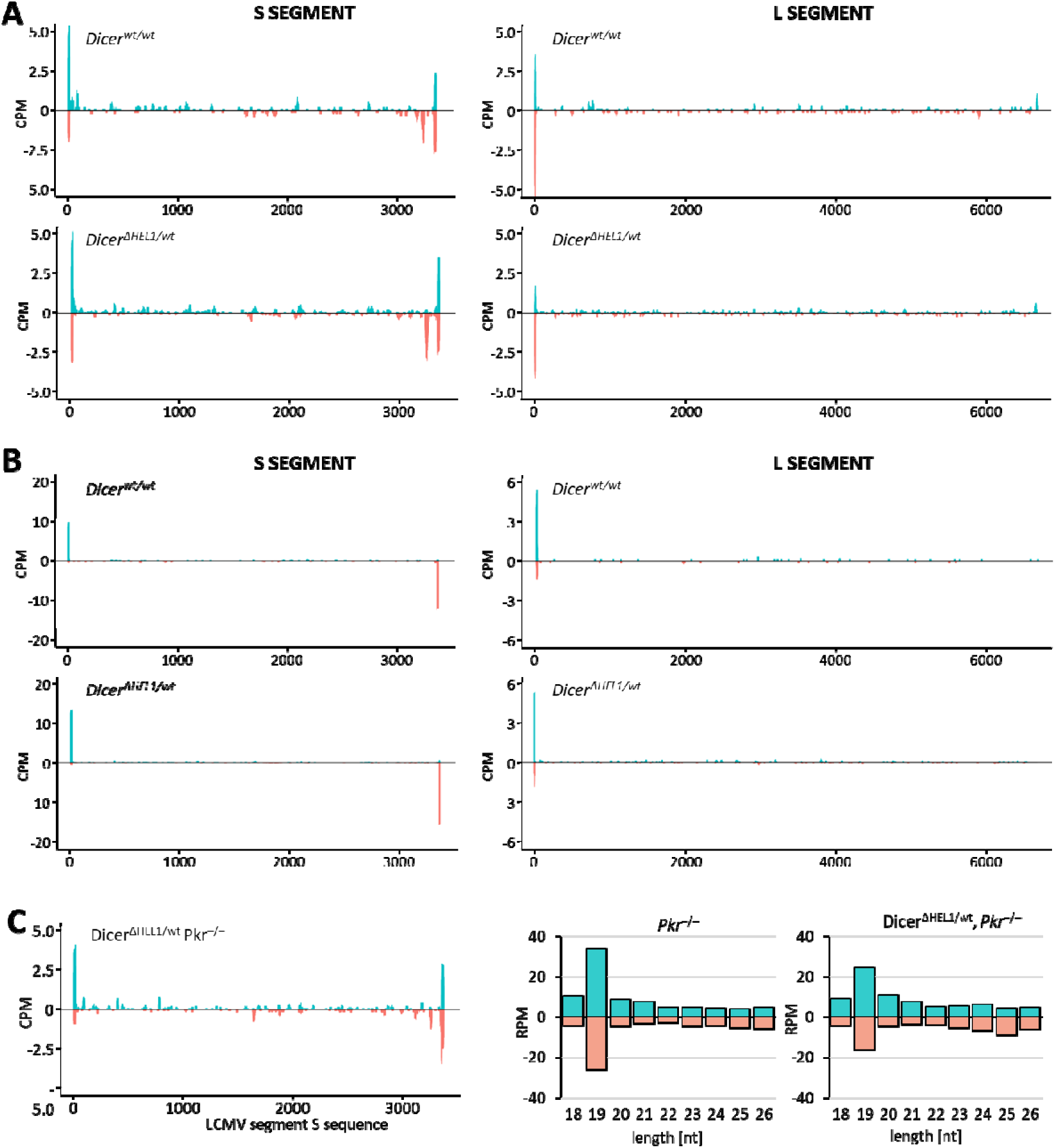
LCMV-derived small RNA analysis in spleen. (A) Coverage plots showing 21-23 nt reads from infected spleen mapped onto S and L segments of the LCMV genome. (B) Coverage plots showing 19 nt reads from infected spleen mapped onto S and L segments of the LCMV genome. (C) Analysis of small RNAs from infected spleen of a *Dicer*^Δ*HEL1/wt*^ *Pkr*^-/-^ mouse, which was infected with 2x10^5^ PFU of LCMV Arm and collected at 3 dpi. Left is a coverage plot for 21-23 nt small RNAs mapping to the S segment. Right is shown analysis distribution of lengths of small RNAs derived from viral RNA in a spleen of infected *Pkr*^-/-^ and *Dicer*^Δ*HEL1/wt*^ *Pkr*^-/-^ mice at 3 dpi. One spleen was sequenced and analyzed for each genotype.

**Figure S4.**
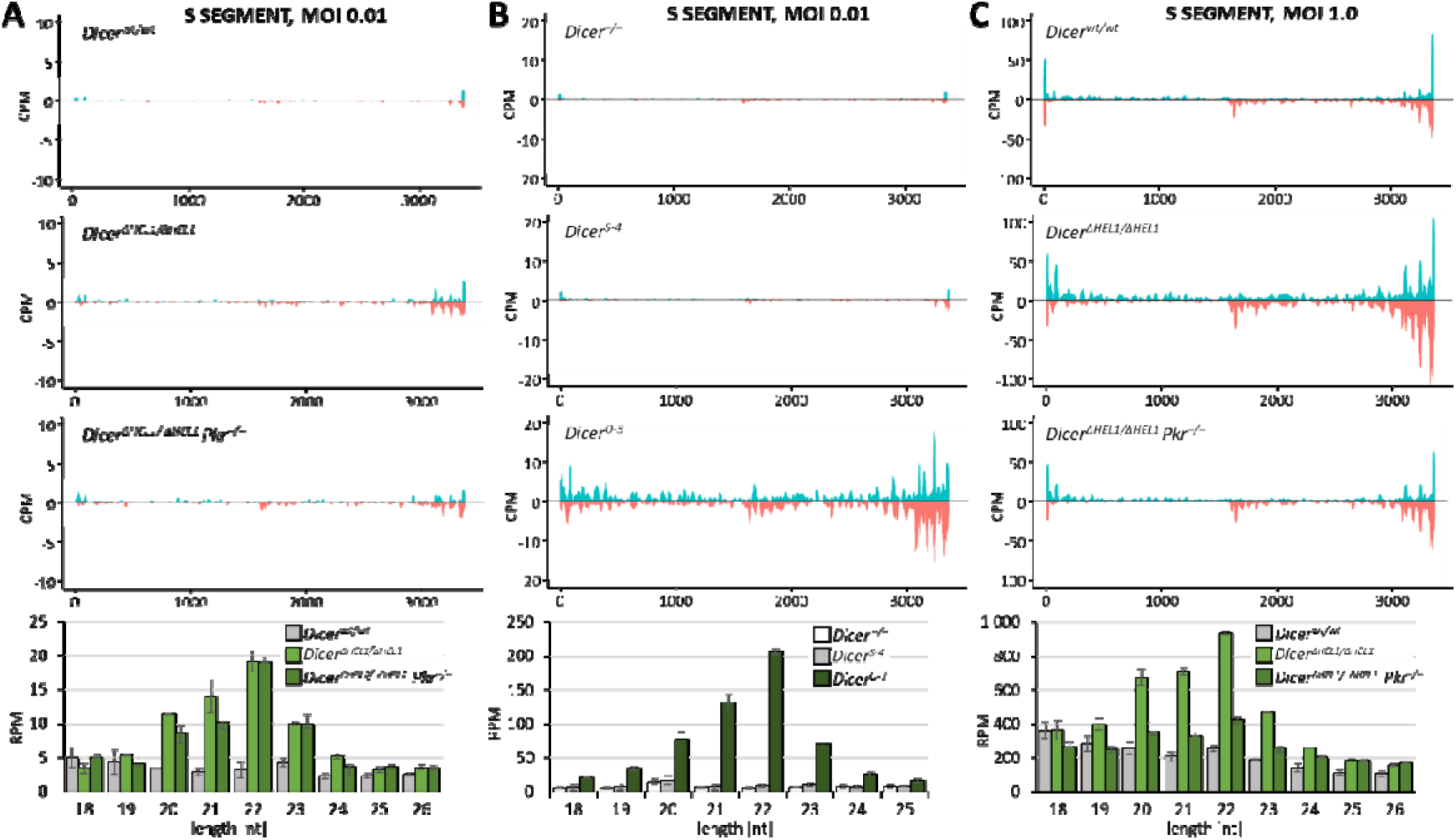
Supporting data for induction of antiviral RNAi in ESCs expressing truncated Dicer. (A) 21-23 nt LCMV abundance in *Dicer*^Δ*HEL1*^ ESCs. Coverage plots show 21-23 nt reads from infected ESCs (MOI 0.01) mapped onto the S segment of the LCMV genome. From above: the parental ESCs line (*Dicer^wt/wt^*), *Dicer*^Δ*HEL1/wt*^,and *Dicer*^Δ*HEL1/wt*^ *Pkr*^-/-^ (the same cell were studied in (Buccheri et al., 2024), Fig. 3). The analysis was performed in a duplicate infection, coverage plots display combined data. The histogram below coverage plots shows analysis of size distribution of small RNAs. Error bars = range of values. (B) 21-23 nt LCMV abundance in *Dicer^O^* ESCs. Coverage plots show 21-23 nt reads from infected ESCs (MOI 0.01) mapped onto the S segment of the LCMV genome. From above: *Dicer^−^*^/*−*^ (Murchison et al., 2005), *Dicer^S-4^* and *Dicer^O-3^* (stable ESC lines where Dicer expression in *Dicer^−^*^/*−*^ was rescued with the full-length Dicer and Dicer^O^, respectively (Flemr et al., 2013)). The analysis was performed in a duplicate infection, coverage plots display combined data. The histogram below coverage plots shows analysis of size distribution of small RNAs. Error bars = range of values. (C) Same as A but with MOI 1.0.

**Table S1.**
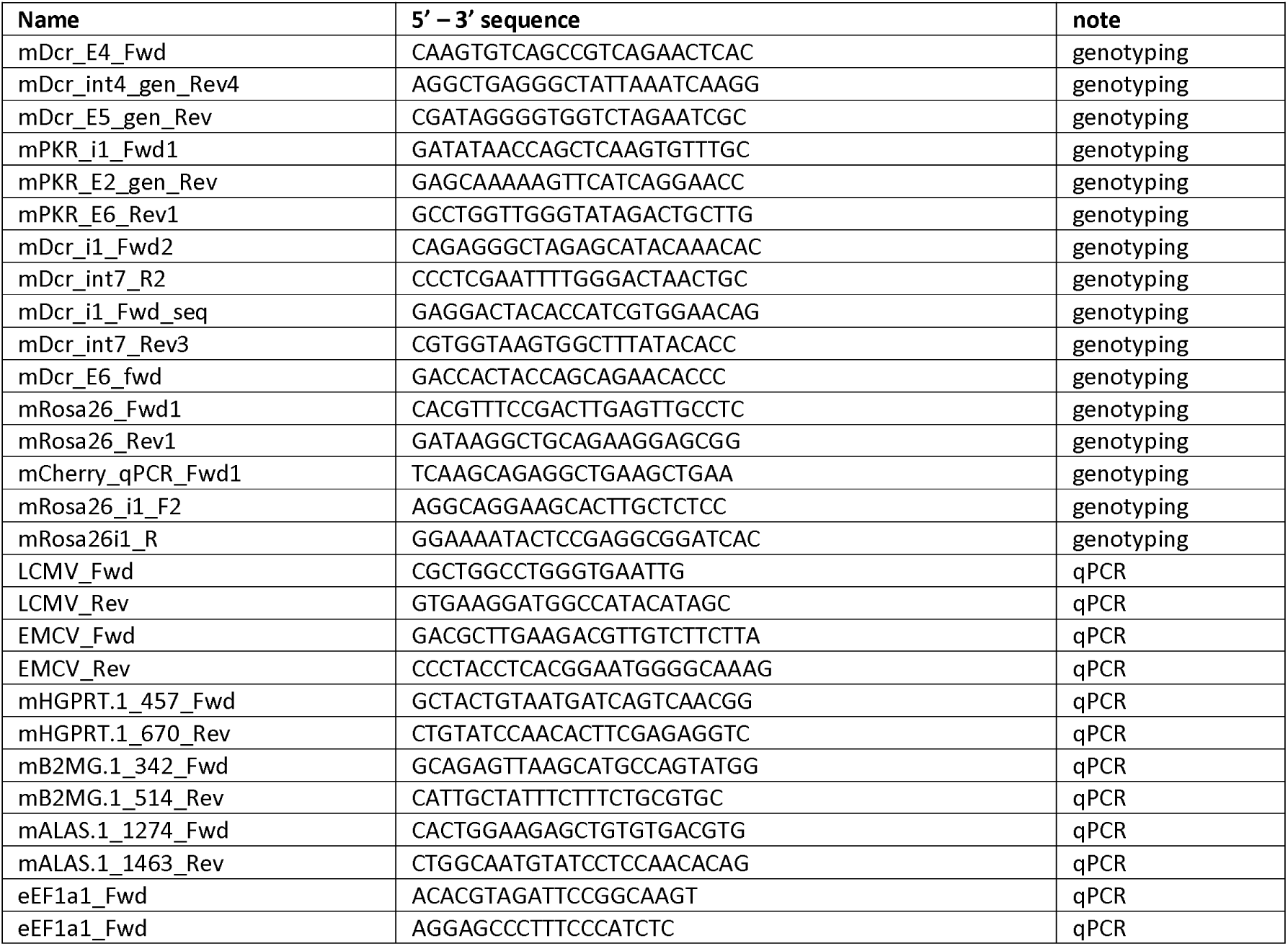
Primers.

**Table S2.** Sequencing libraries deposited in GEO.

| genotype | GEO library name | note | virus | age | organ |
| --- | --- | --- | --- | --- | --- |
| *Dicer*<sup>wt/wt</sup> | spleen_WT_CVB3_B255 | Fig. 1G & S1B | CVB3 | juvenile (4 weeks) | spleen |
| *Dicer*<sup>wt/wt</sup> | spleen_WT_CVB3_S33 | Fig. 1G & S1B | CVB3 | juvenile (4 weeks) | spleen |
| *Dicer*<sup>ΔHEL1/wt</sup> | spleen_DicerDHEL1het_CVB3_S26 | Fig. 1F, G & S1B | CVB3 | juvenile (4 weeks) | spleen |
| *Dicer*<sup>ΔHEL1/wt</sup> | spleen_DicerDHEL1het_CVB3_S25 | Fig. 1F, G & S1B | CVB3 | juvenile (4 weeks) | spleen |
| *Dicer*<sup>wt/wt</sup> | heart_WT_EMCV_46900 | Fig. 2B-D & S1C | EMCV | young adult (6 weeks) | heart |
| *Dicer*<sup>wt/wt</sup> | heart_WT_EMCV_46906 | Fig. 2B-D & S1C | EMCV | young adult (6 weeks) | heart |
| *Dicer*<sup>ΔHEL1/wt</sup> | heart_DHEL1het_EMCV_46908 | Fig. 2B-D & S1C | EMCV | young adult (6 weeks) | heart |
| *Dicer*<sup>ΔHEL1/wt</sup> | heart_DHEL1het_EMCV_46907 | Fig. 2B-D & S1C | EMCV | young adult (6 weeks) | heart |
| *Dicer*<sup>wt/wt</sup> | brain_WT_TBEBV_4 | Fig. 3D-F, S2A & S2C | TBEV | young adult (6-7 weeks) | brain |
| *Dicer*<sup>wt/wt</sup> | brain_WT_TBEBV_5 | Fig. 3D-F, S2A & S2C | TBEV | young adult (6-7 weeks) | brain |
| *Dicer*<sup>wt/wt</sup> | brain_WT_TBEBV_6 | Fig. 3D-F, S2A & S2C | TBEV | young adult (6-7 weeks) | brain |
| *Dicer*<sup>ΔHEL1/wt</sup> | brain_DicerDHEL1het_TBEBV_1 | Fig. 3D-F, S2A & S2C | TBEV | young adult (6-7 weeks) | brain |
| *Dicer*<sup>ΔHEL1/wt</sup> | brain_DicerDHEL1het_TBEBV_2 | Fig. 3D-F, S2A & S2C | TBEV | young adult (6-7 weeks) | brain |
| *Dicer*<sup>ΔHEL1/wt</sup> | brain_DicerDHEL1het_TBEBV_3 | Fig. 3D-F, S2A & S2C | TBEV | young adult (6-7 weeks) | brain |
| *Dicer*<sup>wt/wt</sup>, *Pkr*<sup>-/-</sup> | brain_PKRnull_TBEBV_3dpi | Fig. 3J & S1B | TBEV | young adult (6-7 weeks) | brain |
| *Dicer*<sup>wt/wt</sup>, *Pkr*<sup>-/-</sup> | brain_PKRnull_TBEBV_6dpi | Fig. 3J & S1B | TBEV | young adult (6-7 weeks) | brain |
| *Dicer*<sup>wt/wt</sup>, *Pkr*<sup>-/-</sup> | brain_PKRnull_TBEBV_10dpi | Fig. 3J & S1B | TBEV | young adult (6-7 weeks) | brain |
| *Dicer*<sup>ΔHEL1/wt</sup>, *Pkr*<sup>-/-</sup> | brain_DicerDHEL1het_PKRnull_TBEBV_3dpi | Fig. 3J & S1B | TBEV | young adult (6-7 weeks) | brain |
| *Dicer*<sup>ΔHEL1/wt</sup>, *Pkr*<sup>-/-</sup> | brain_DicerDHEL1het_PKRnull_TBEBV_6dpi | Fig. 3J & S1B | TBEV | young adult (6-7 weeks) | brain |
| *Dicer*<sup>ΔHEL1/wt</sup>, *Pkr*<sup>-/-</sup> | brain_DicerDHEL1het_PKRnull_TBEBV_10dpi | Fig. 3J & S1B | TBEV | young adult (6-7 weeks) | brain |
| *Dicer*<sup>wt/wt</sup> | spleen_WT_LCMV_28Y35226 | Fig. 4C & S3 | LCMV | adult (12 weeks) | spleen |
| *Dicer*<sup>wt/wt</sup> | spleen_WT_LCMV_28Y35286 | Fig. 4C & S3 | LCMV | adult (12 weeks) | spleen |
| *Dicer*<sup>ΔHEL1/wt</sup> | spleen_DicerDHEL1het_LCMV_28Y35225 | Fig. 4B-D & S3 | LCMV | adult (12 weeks) | spleen |
| *Dicer*<sup>ΔHEL1/wt</sup> | spleen_DicerDHEL1het_LCMV_28Y35291 | Fig. 4B-D & S3 | LCMV | adult (12 weeks) | spleen |
| *Dicer*<sup>tg(O-HA)/tg(O-HA)</sup> | spleen_DicerTG_LCMV_28Y35336 | Fig. 6E | LCMV | adult (12 weeks) | spleen |
| *Dicer*<sup>tg(O-HA)/tg(O-HA)</sup> | spleen_DicerTG_LCMV_28Y35338 | Fig. 6E | LCMV | adult (12 weeks) | spleen |
| *Dicer*<sup>wt/wt</sup>, *Pkr*<sup>-/-</sup> | spleen_PKRnull_LCMV_3dpi | Fig. 4C & S3 | LCMV | adult (12 weeks) | spleen |
| *Dicer*<sup>ΔHEL1/wt</sup>, *Pkr*<sup>-/-</sup> | spleen_DicerDHEL1het_PKRnull_LCMV_3dpi | Fig. 4C & S3 | LCMV | adult (12 weeks) | spleen |
| *Dicer*<sup>wt/wt</sup> | mESC_WT_LCMV_MOI_0.01_7 | Fig. 5E, S4A | LCMV | cell line | ESC |
| *Dicer*<sup>wt/wt</sup> | mESC_WT_LCMV_MOI_0.01_8 | Fig. 5E, S4A | LCMV | cell line | ESC |
| *Dicer*<sup>ΔHEL1/ΔHEL1</sup> | mESC_DicerDHEL1_LCMV_MOI_0.01_9 | Fig. 5E, S4A | LCMV | cell line | ESC |
| *Dicer*<sup>ΔHEL1/ΔHEL1</sup> | mESC_DicerDHEL1_LCMV_MOI_0.01_10 | Fig. 5E, S4A | LCMV | cell line | ESC |
| *Dicer*<sup>ΔHEL1/ΔHEL1</sup>, *Pkr*<sup>-/-</sup> | mESC_DicerDHEL1_PKRnull_LCMV_MOI_0.01_11 | Fig. S4A | LCMV | cell line | ESC |
| *Dicer*<sup>ΔHEL1/ΔHEL1</sup>, *Pkr*<sup>-/-</sup> | mESC_DicerDHEL1_PKRnull_LCMV_MOI_0.01_12 | Fig. S4A | LCMV | cell line | ESC |
| *Dicer*<sup>wt/wt</sup> | mESC_WT_LCMV_MOI_1_15 | Fig. S4C | LCMV | cell line | ESC |
| *Dicer*<sup>wt/wt</sup> | mESC_WT_LCMV_MOI_1_16 | Fig. S4C | LCMV | cell line | ESC |
| *Dicer*<sup>ΔHEL1/ΔHEL1</sup> | mESC_DicerDHEL1_LCMV_MOI_1_19 | Fig. S4C | LCMV | cell line | ESC |
| *Dicer*<sup>ΔHEL1/ΔHEL1</sup> | mESC_DicerDHEL1_LCMV_MOI_1_20 | Fig. S4C | LCMV | cell line | ESC |
| *Dicer*<sup>ΔHEL1/ΔHEL1</sup>, *Pkr*<sup>-/-</sup> | mESC_DicerDHEL1_PKRnull_LCMV_MOI_1_23 | Fig. S4C | LCMV | cell line | ESC |
| *Dicer*<sup>ΔHEL1/ΔHEL1</sup>, *Pkr*<sup>-/-</sup> | mESC_DicerDHEL1_PKRnull_LCMV_MOI_1_24 | Fig. S4C | LCMV | cell line | ESC |
| *Dicer*<sup>-/-</sup> | mESC_DicerKO_LCMV_MOI_0.01_1 | Fig. S4B | LCMV | cell line | ESC |
| *Dicer*<sup>-/-</sup> | mESC_DicerKO_LCMV_MOI_0.01_2 | Fig. S4B | LCMV | cell line | ESC |
| *Dicer*<sup>O-3</sup> | mESC_DicerO3_LCMV_MOI_0.01_1 | Fig. 5D, 5E, S4B | LCMV | cell line | ESC |
| *Dicer*<sup>O-3</sup> | mESC_DicerO3_LCMV_MOI_0.01_2 | Fig. 5D, 5E, S4B | LCMV | cell line | ESC |
| *Dicer*<sup>S-4</sup> | mESC_DicerS4_LCMV_MOI_0.01_1 | Fig. S4B | LCMV | cell line | ESC |
| *Dicer*<sup>S-4</sup> | mESC_DicerS4_LCMV_MOI_0.01_2 | Fig. S4B | LCMV | cell line | ESC |
| *Dicer*<sup>wt/wt</sup> | mESC_WT_LCMV_MOI_0.01_S9 | Fig. 5G | LCMV | cell line | ESC |
| *Dicer*<sup>wt/wt</sup> | mESC_WT_LCMV_MOI_0.01_S10 | Fig. 5G | LCMV | cell line | ESC |
| *Dicer*<sup>ΔHEL1/ΔHEL1</sup> | mESC_DicerDHEL1_LCMV_MOI_0.01_S11 | Fig. 5G | LCMV | cell line | ESC |
| *Dicer*<sup>ΔHEL1/ΔHEL1</sup> | mESC_DicerDHEL1_LCMV_MOI_0.01_S12 | Fig. 5G | LCMV | cell line | ESC |
| *Dicer*<sup>O-3</sup> | mESC_DicerO3_LCMV_MOI_0.01_S15 | Fig. 5G | LCMV | cell line | ESC |
| *Dicer*<sup>O-3</sup> | mESC_DicerO3_LCMV_MOI_0.01_S16 | Fig. 5G | LCMV | cell line | ESC |
| *Dicer*<sup>wt/wt</sup> | mESC_WT_MosIR_S1 | Fig. 5H | MosIR | cell line | ESC |
| *Dicer*<sup>wt/wt</sup> | mESC_WT_MosIR_S2 | Fig. 5H | MosIR | cell line | ESC |
| *Dicer*<sup>ΔHEL1/ΔHEL1</sup> | mESC_DicerDHEL1_MosIR_S3 | Fig. 5H | MosIR | cell line | ESC |
| *Dicer*<sup>ΔHEL1/ΔHEL1</sup> | mESC_DicerDHEL1_MosIR_S4 | Fig. 5H | MosIR | cell line | ESC |
| *Dicer*<sup>O-3</sup> | mESC_DicerO3_MosIR_S7 | Fig. 5H | MosIR | cell line | ESC |
| *Dicer*<sup>O-3</sup> | mESC_DicerO3_MosIR_S8 | Fig. 5H | MosIR | cell line | ESC |
| *Dicer*<sup>wt/wt</sup> | mESC_WT_LCMV_MOI_0.01_TrapR_S9 | Fig. 5G | LCMV | cell line | ESC |
| *Dicer*<sup>wt/wt</sup> | mESC_WT_LCMV_MOI_0.01_TrapR_S10 | Fig. 5G | LCMV | cell line | ESC |
| *Dicer*<sup>ΔHEL1/ΔHEL1</sup> | mESC_DicerDHEL1_LCMV_MOI_0.01_TrapR_S11 | Fig. 5G | LCMV | cell line | ESC |
| *Dicer*<sup>ΔHEL1/ΔHEL1</sup> | mESC_DicerDHEL1_LCMV_MOI_0.01_TrapR_S12 | Fig. 5G | LCMV | cell line | ESC |
| *Dicer*<sup>O-3</sup> | mESC_DicerO3_LCMV_MOI_0.01_TrapR_S15 | Fig. 5G | LCMV | cell line | ESC |
| *Dicer*<sup>O-3</sup> | mESC_DicerO3_LCMV_MOI_0.01_TrapR_S16 | Fig. 5G | LCMV | cell line | ESC |
| *Dicer*<sup>wt/wt</sup> | mESC_WT_MosIR_TrapR_S1 | Fig. 5H | MosIR | cell line | ESC |
| *Dicer*<sup>wt/wt</sup> | mESC_WT_MosIR_TrapR_S2 | Fig. 5H | MosIR | cell line | ESC |
| *Dicer*<sup>ΔHEL1/ΔHEL1</sup> | mESC_DicerDHEL1_MosIR_TrapR_S3 | Fig. 5H | MosIR | cell line | ESC |
| *Dicer*<sup>ΔHEL1/ΔHEL1</sup> | mESC_DicerDHEL1_MosIR_TrapR_S4 | Fig. 5H | MosIR | cell line | ESC |
| *Dicer*<sup>O-3</sup> | mESC_DicerO3_MosIR_TrapR_S7 | Fig. 5H | MosIR | cell line | ESC |
| *Dicer*<sup>O-3</sup> | mESC_DicerO3_MosIR_TrapR_S8 | Fig. 5H | MosIR | cell line | ESC |

## References

1. Aderounmu, A.M., Aruscavage, P.J., Kolaczkowski, B., and Bass, B.L. (2023). Ancestral protein reconstruction reveals evolutionary events governing variation in Dicer helicase function. Elife 12.

2. Agudelo, M., Palus, M., Keeffe, J.R., Bianchini, F., Svoboda, P., Salat, J., Peace, A., Gazumyan, A., Cipolla, M., Kapoor, T., et al. (2021). Broad and potent neutralizing human antibodies to tick-borne flaviviruses protect mice from disease. J Exp Med 218.

3. Ameres, S.L., Horwich, M.D., Hung, J.H., Xu, J., Ghildiyal, M., Weng, Z., and Zamore, P.D. (2010). Target RNA-directed trimming and tailing of small silencing RNAs. Science 328, 1534–1539.

4. Baldaccini, M., Gaucherand, L., Chane-Woon-Ming, B., Messmer, M., Gucciardi, F., and Pfeffer, S. (2024). The helicase domain of human Dicer prevents RNAi-independent activation of antiviral and inflammatory pathways. EMBO J 43, 806–835.

5. Bartel, D.P. (2018). Metazoan MicroRNAs. Cell 173, 20–51.

6. Bernstein, E., Kim, S.Y., Carmell, M.A., Murchison, E.P., Alcorn, H., Li, M.Z., Mills, A.A., Elledge, S.J., Anderson, K.V., and Hannon, G.J. (2003). Dicer is essential for mouse development. Nat Genet 35, 215–217.

7. Bogerd, H.P., Skalsky, R.L., Kennedy, E.M., Furuse, Y., Whisnant, A.W., Flores, O., Schultz, K.L., Putnam, N., Barrows, N.J., Sherry, B., et al. (2014). Replication of many human viruses is refractory to inhibition by endogenous cellular microRNAs. J Virol 88, 8065–8076.

8. Buccheri, V., Pasulka, J., Malik, R., Loubalova, Z., Taborska, E., Horvat, F., Roos Kulmann, M.I., Jenickova, I., Prochazka, J., Sedlacek, R., et al. (2024). Functional canonical RNAi in mice expressing a truncated Dicer isoform and long dsRNA. EMBO Reports in press.

9. Bushnell, B. (2015). BBMap short read aligner and other bioinformatics tools.

10. Carocci, M., and Bakkali-Kassimi, L. (2012). The encephalomyocarditis virus. Virulence 3, 351–367.

11. Cullen, B.R., Cherry, S., and tenOever, B.R. (2013). Is RNA interference a physiologically relevant innate antiviral immune response in mammals? Cell Host Microbe 14, 374–378.

12. De Madrid, A.T., and Porterfield, J.S. (1969). A simple micro-culture method for the study of group B arboviruses. Bull World Health Organ 40, 113–121.

13. Demeter, T., Vaskovicova, M., Malik, R., Horvat, F., Pasulka, J., Svobodova, E., Flemr, M., and Svoboda, P. (2019). Main constraints for RNAi induced by expressed long dsRNA in mouse cells. Life Sci Alliance 2.

14. Dobin, A., Davis, C.A., Schlesinger, F., Drenkow, J., Zaleski, C., Jha, S., Batut, P., Chaisson, M., and Gingeras, T.R. (2013). STAR: ultrafast universal RNA-seq aligner. Bioinformatics 29, 15–21.

15. Fang, Y., Liu, Z., Qiu, Y., Kong, J., Fu, Y., Liu, Y., Wang, C., Quan, J., Wang, Q., Xu, W., et al. (2021). Inhibition of viral suppressor of RNAi proteins by designer peptides protects from enteroviral infection in vivo. Immunity 54, 2231–2244 e2236.

16. Felix, M.A., Ashe, A., Piffaretti, J., Wu, G., Nuez, I., Belicard, T., Jiang, Y., Zhao, G., Franz, C.J., Goldstein, L.D., et al. (2011). Natural and experimental infection of Caenorhabditis nematodes by novel viruses related to nodaviruses. PLoS Biol 9, e1000586.

17. Fire, A., Xu, S., Montgomery, M.K., Kostas, S.A., Driver, S.E., and Mello, C.C. (1998). Potent and specific genetic interference by double-stranded RNA in Caenorhabditis elegans. Nature 391, 806–811.

18. Flemr, M., Malik, R., Franke, V., Nejepinska, J., Sedlacek, R., Vlahovicek, K., and Svoboda, P. (2013). A retrotransposon-driven Dicer isoform directs endogenous small interfering RNA production in mouse oocytes. Cell 155, 807–816.

19. Frankish, A., Diekhans, M., Ferreira, A.M., Johnson, R., Jungreis, I., Loveland, J., Mudge, J.M., Sisu, C., Wright, J., Armstrong, J., et al. (2019). GENCODE reference annotation for the human and mouse genomes. Nucleic Acids Res 47, D766–D773.

20. Garmaroudi, F.S., Marchant, D., Hendry, R., Luo, H., Yang, D., Ye, X., Shi, J., and McManus, B.M. (2015). Coxsackievirus B3 replication and pathogenesis. Future Microbiol 10, 629–653.

21. Girardi, E., Chane-Woon-Ming, B., Messmer, M., Kaukinen, P., and Pfeffer, S. (2013). Identification of RNase L-dependent, 3’-end-modified, viral small RNAs in Sindbis virus-infected mammalian cells. mBio 4, e00698–00613.

22. Girardi, E., Lefevre, M., Chane-Woon-Ming, B., Paro, S., Claydon, B., Imler, J.L., Meignin, C., and Pfeffer, S. (2015). Cross-species comparative analysis of Dicer proteins during Sindbis virus infection. Sci Rep 5, 10693.

23. Grentzinger, T., Oberlin, S., Schott, G., Handler, D., Svozil, J., Barragan-Borrero, V., Humbert, A., Duharcourt, S., Brennecke, J., and Voinnet, O. (2020). A universal method for the rapid isolation of all known classes of functional silencing small RNAs. Nucleic Acids Res 48, e79.

24. Han, Q., Chen, G., Wang, J., Jee, D., Li, W.X., Lai, E.C., and Ding, S.W. (2020). Mechanism and Function of Antiviral RNA Interference in Mice. mBio 11.

25. Hierholzer, E., and Killington, R.A. (1996). Virus isolation and quantitation. In Virology methods manual, B.W. Mahy, and H.O. Kangro, eds. (San Diego: Academic Press), pp. 25-46.

26. Horkova, V., Drobek, A., Paprckova, D., Niederlova, V., Prasai, A., Uleri, V., Glatzova, D., Kraller, M., Cesnekova, M., Janusova, S., et al. (2023). Unique roles of co-receptor-bound LCK in helper and cytotoxic T cells. Nat Immunol 24, 174–185.

27. Chakravarthy, S., Sternberg, S.H., Kellenberger, C.A., and Doudna, J.A. (2010). Substrate-specific kinetics of Dicer-catalyzed RNA processing. J Mol Biol 404, 392–402.

28. Cheloufi, S., Dos Santos, C.O., Chong, M.M., and Hannon, G.J. (2010). A dicer-independent miRNA biogenesis pathway that requires Ago catalysis. Nature 465, 584–589.

29. Kakumani, P.K., Ponia, S.S., S, R.K., Sood, V., Chinnappan, M., Banerjea, A.C., Medigeshi, G.R., Malhotra, P., Mukherjee, S.K., and Bhatnagar, R.K. (2013). Role of RNA interference (RNAi) in dengue virus replication and identification of NS4B as an RNAi suppressor. J Virol 87, 8870–8883.

30. Kennedy, E.M., Whisnant, A.W., Kornepati, A.V., Marshall, J.B., Bogerd, H.P., and Cullen, B.R. (2015). Production of functional small interfering RNAs by an amino-terminal deletion mutant of human Dicer. Proc Natl Acad Sci U S A.

31. Ketting, R.F. (2011). The many faces of RNAi. Dev Cell 20, 148–161.

32. Kozomara, A., Birgaoanu, M., and Griffiths-Jones, S. (2019). miRBase: from microRNA sequences to function. Nucleic Acids Res 47, D155–D162.

33. Laposova, K., Pastorekova, S., and Tomaskova, J. (2013). Lymphocytic choriomeningitis virus: invisible but not innocent. Acta Virol 57, 160–170.

34. Li, Y., Basavappa, M., Lu, J., Dong, S., Cronkite, D.A., Prior, J.T., Reinecker, H.C., Hertzog, P., Han, Y., Li, W.X., et al. (2016). Induction and suppression of antiviral RNA interference by influenza A virus in mammalian cells. Nat Microbiol 2, 16250.

35. Li, Y., Lu, J., Han, Y., Fan, X., and Ding, S.W. (2013). RNA interference functions as an antiviral immunity mechanism in mammals. Science 342, 231–234.

36. Liao, Y., Smyth, G.K., and Shi, W. (2014). featureCounts: an efficient general purpose program for assigning sequence reads to genomic features. Bioinformatics 30, 923–930.

37. Lindberg, A.M., Stalhandske, P.O., and Pettersson, U. (1987). Genome of coxsackievirus B3. Virology 156, 50–63.

38. Liu, J.D., Carmell, M.A., Rivas, F.V., Marsden, C.G., Thomson, J.M., Song, J.J., Hammond, S.M., Joshua-Tor, L., and Hannon, G.J. (2004). Argonaute2 is the catalytic engine of mammalian RNAi. Science 305, 1437–1441.

39. Liu, Z., Wang, J., Cheng, H., Ke, X., Sun, L., Zhang, Q.C., and Wang, H.W. (2018). Cryo-EM Structure of Human Dicer and Its Complexes with a Pre-miRNA Substrate. Cell 173, 1191–1203 e1112.

40. Love, M.I., Huber, W., and Anders, S. (2014). Moderated estimation of fold change and dispersion for RNA-seq data with DESeq2. Genome Biol 15, 550.

41. Lu, R., Maduro, M., Li, F., Li, H.W., Broitman-Maduro, G., Li, W.X., and Ding, S.W. (2005). Animal virus replication and RNAi-mediated antiviral silencing in Caenorhabditis elegans. Nature 436, 1040–1043.

42. Ma, E., MacRae, I.J., Kirsch, J.F., and Doudna, J.A. (2008). Autoinhibition of human dicer by its internal helicase domain. J Mol Biol 380, 237–243.

43. Maillard, P.V., Ciaudo, C., Marchais, A., Li, Y., Jay, F., Ding, S.W., and Voinnet, O. (2013). Antiviral RNA interference in mammalian cells. Science 342, 235–238.

44. Martin, M. (2011). Cutadapt removes adapter sequences from high-throughput sequencing reads. 2011 *17*, 3.

45. Meister, G., Landthaler, M., Patkaniowska, A., Dorsett, Y., Teng, G., and Tuschl, T. (2004). Human Argonaute2 mediates RNA cleavage targeted by miRNAs and siRNAs. Mol Cell 15, 185–197.

46. Miyazaki, J., Takaki, S., Araki, K., Tashiro, F., Tominaga, A., Takatsu, K., and Yamamura, K. (1989). Expression vector system based on the chicken beta-actin promoter directs efficient production of interleukin-5. Gene 79, 269–277.

47. Montavon, T.C., Baldaccini, M., Lefevre, M., Girardi, E., Chane-Woon-Ming, B., Messmer, M., Hammann, P., Chicher, J., and Pfeffer, S. (2021). Human DICER helicase domain recruits PKR and modulates its antiviral activity. PLoS Pathog 17, e1009549.

48. Murchison, E.P., Partridge, J.F., Tam, O.H., Cheloufi, S., and Hannon, G.J. (2005). Characterization of Dicer-deficient murine embryonic stem cells. Proc Natl Acad Sci U S A 102, 12135–12140.

49. Nejepinska, J., Malik, R., Filkowski, J., Flemr, M., Filipowicz, W., and Svoboda, P. (2012). dsRNA expression in the mouse elicits RNAi in oocytes and low adenosine deamination in somatic cells. Nucleic Acids Res 40, 399–413.

50. Okabe, M., Ikawa, M., Kominami, K., Nakanishi, T., and Nishimune, Y. (1997). ’Green mice’ as a source of ubiquitous green cells. FEBS letters 407, 313–319.

51. Parameswaran, P., Sklan, E., Wilkins, C., Burgon, T., Samuel, M.A., Lu, R., Ansel, K.M., Heissmeyer, V., Einav, S., Jackson, W., et al. (2010). Six RNA viruses and forty-one hosts: viral small RNAs and modulation of small RNA repertoires in vertebrate and invertebrate systems. PLoS Pathog 6, e1000764.

52. Poirier, E.Z., Buck, M.D., Chakravarty, P., Carvalho, J., Frederico, B., Cardoso, A., Healy, L., Ulferts, R., Beale, R., and Reis e Sousa, C. (2021). An isoform of Dicer protects mammalian stem cells against multiple RNA viruses. Science 373, 231–236.

53. Pokorna Formanova, P., Palus, M., Salat, J., Honig, V., Stefanik, M., Svoboda, P., and Ruzek, D. (2019). Changes in cytokine and chemokine profiles in mouse serum and brain, and in human neural cells, upon tick-borne encephalitis virus infection. J Neuroinflammation 16, 205.

54. Qiu, Y., Xu, Y., Zhang, Y., Zhou, H., Deng, Y.Q., Li, X.F., Miao, M., Zhang, Q., Zhong, B., Hu, Y., et al. (2017). Human Virus-Derived Small RNAs Can Confer Antiviral Immunity in Mammals. Immunity 46, 992–1004 e1005.

55. Qiu, Y., Xu, Y.P., Wang, M., Miao, M., Zhou, H., Xu, J., Kong, J., Zheng, D., Li, R.T., Zhang, R.R., et al. (2020). Flavivirus induces and antagonizes antiviral RNA interference in both mammals and mosquitoes. Sci Adv 6, eaax7989.

56. Saleh, M.C., Tassetto, M., van Rij, R.P., Goic, B., Gausson, V., Berry, B., Jacquier, C., Antoniewski, C., and Andino, R. (2009). Antiviral immunity in Drosophila requires systemic RNA interference spread. Nature 458, 346–350.

57. Sanchez, A.B., Perez, M., Cornu, T., and de la Torre, J.C. (2005). RNA interference-mediated virus clearance from cells both acutely and chronically infected with the prototypic arenavirus lymphocytic choriomeningitis virus. J Virol 79, 11071–11081.

58. Sarkies, P., Ashe, A., Le Pen, J., McKie, M.A., and Miska, E.A. (2013). Competition between virus-derived and endogenous small RNAs regulates gene expression in Caenorhabditis elegans. Genome research 23, 1258–1270.

59. Seo, G.J., Kincaid, R.P., Phanaksri, T., Burke, J.M., Pare, J.M., Cox, J.E., Hsiang, T.Y., Krug, R.M., and Sullivan, C.S. (2013). Reciprocal inhibition between intracellular antiviral signaling and the RNAi machinery in mammalian cells. Cell Host Microbe 14, 435–445.

60. Schnettler, E., Tykalova, H., Watson, M., Sharma, M., Sterken, M.G., Obbard, D.J., Lewis, S.H., McFarlane, M., Bell-Sakyi, L., Barry, G., et al. (2014). Induction and suppression of tick cell antiviral RNAi responses by tick-borne flaviviruses. Nucleic Acids Res 42, 9436–9446.

61. Schott, D.H., Cureton, D.K., Whelan, S.P., and Hunter, C.P. (2005). An antiviral role for the RNA interference machinery in Caenorhabditis elegans. Proc Natl Acad Sci U S A 102, 18420–18424.

62. Schuster, S., Overheul, G.J., Bauer, L., van Kuppeveld, F.J.M., and van Rij, R.P. (2019). No evidence for viral small RNA production and antiviral function of Argonaute 2 in human cells. Sci Rep 9, 13752.

63. Schuster, S., Tholen, L.E., Overheul, G.J., van Kuppeveld, F.J.M., and van Rij, R.P. (2017). Deletion of Cytoplasmic Double-Stranded RNA Sensors Does Not Uncover Viral Small Interfering RNA Production in Human Cells. mSphere 2.

64. Sinkkonen, L., Hugenschmidt, T., Filipowicz, W., and Svoboda, P. (2010). Dicer is associated with ribosomal DNA chromatin in mammalian cells. PLoS One 5, e12175.

65. Song, J.J., Smith, S.K., Hannon, G.J., and Joshua-Tor, L. (2004). Crystal structure of Argonaute and its implications for RISC slicer activity. Science 305, 1434–1437.

66. Srivastava, B., Blazejewska, P., Hessmann, M., Bruder, D., Geffers, R., Mauel, S., Gruber, A.D., and Schughart, K. (2009). Host genetic background strongly influences the response to influenza a virus infections. PLoS One 4, e4857.

67. Svoboda, P., Stein, P., and Schultz, R.M. (2001). RNAi in mouse oocytes and preimplantation embryos: Effectiveness of hairpin dsRNA. Biochemical and Biophysical Research Communications 287, 1099–1104.

68. Taborska, E., Pasulka, J., Malik, R., Horvat, F., Jenickova, I., Jelic Matosevic, Z., and Svoboda, P. (2019). Restricted and non-essential redundancy of RNAi and piRNA pathways in mouse oocytes. PLoS Genet 15, e1008261.

69. Taborska, E.e.a. (2024). Activated RNAi does not rescue piRNA pathway deficiency in testes. manuscript.

70. Takahashi, T., Nakano, Y., Onomoto, K., Yoneyama, M., and Ui-Tei, K. (2018). Virus Sensor RIG-I Represses RNA Interference by Interacting with TRBP through LGP2 in Mammalian Cells. Genes (Basel) 9.

71. Tassetto, M., Kunitomi, M., and Andino, R. (2017). Circulating Immune Cells Mediate a Systemic RNAi-Based Adaptive Antiviral Response in Drosophila. Cell 169, 314–325 e313.

72. Toni, L.S., Garcia, A.M., Jeffrey, D.A., Jiang, X., Stauffer, B.L., Miyamoto, S.D., and Sucharov, C.C. (2018). Optimization of phenol-chloroform RNA extraction. MethodsX 5, 599–608.

73. van der Veen, A.G., Maillard, P.V., Schmidt, J.M., Lee, S.A., Deddouche-Grass, S., Borg, A., Kjaer, S., Snijders, A.P., and Reis e Sousa, C. (2018). The RIG-I-like receptor LGP2 inhibits Dicer-dependent processing of long double-stranded RNA and blocks RNA interference in mammalian cells. EMBO J 37.

74. Wang, C., Yue, F., and Kuang, S. (2017). Muscle Histology Characterization Using H&E Staining and Muscle Fiber Type Classification Using Immunofluorescence Staining. Bio Protoc 7.

75. Wilkins, C., Dishongh, R., Moore, S.C., Whitt, M.A., Chow, M., and Machaca, K. (2005). RNA interference is an antiviral defence mechanism in Caenorhabditis elegans. Nature 436, 1044–1047.

76. Wilson, R.C., Tambe, A., Kidwell, M.A., Noland, C.L., Schneider, C.P., and Doudna, J.A. (2015). Dicer-TRBP complex formation ensures accurate mammalian microRNA biogenesis. Mol Cell 57, 397–407.

77. Xu, Y.P., Qiu, Y., Zhang, B., Chen, G., Chen, Q., Wang, M., Mo, F., Xu, J., Wu, J., Zhang, R.R., et al. (2019). Zika virus infection induces RNAi-mediated antiviral immunity in human neural progenitors and brain organoids. Cell Res 29, 265–273.

78. Zapletal, D., Kubicek, K., Svoboda, P., and Stefl, R. (2023). Dicer structure and function: conserved and evolving features. EMBO Rep 24, e57215.

79. Zapletal, D., Taborska, E., Pasulka, J., Malik, R., Kubicek, K., Zanova, M., Much, C., Sebesta, M., Buccheri, V., Horvat, F., et al. (2022). Structural and functional basis of mammalian microRNA biogenesis by Dicer. Mol Cell 82, 4064–4079 e4013.

80. Zhang, H., Kolb, F.A., Brondani, V., Billy, E., and Filipowicz, W. (2002). Human Dicer preferentially cleaves dsRNAs at their termini without a requirement for ATP. EMBO J 21, 5875–5885.

81. Zhang, Y., Dai, Y., Wang, J., Xu, Y., Li, Z., Lu, J., Xu, Y., Zhong, J., Ding, S.W., and Li, Y. (2022). Mouse circulating extracellular vesicles contain virus-derived siRNAs active in antiviral immunity. EMBO J 41, e109902.

